# Mapping the chaperonin TRiC/CCT interactome in mouse photoreceptors reveals functional significance for energy metabolism

**DOI:** 10.64898/2026.03.12.711399

**Authors:** Celine Brooks, David Salcedo-Tacuma, Isabella Mascari, Mark Eminhizer, Tuan Ngo, Douglas Kolson, Emily Sechrest, Tongju Guan, Neil Billington, Wen-Tao Deng, David Smith, Jianhai Du, Nikolai Skiba, Maxim Sokolov

## Abstract

The eukaryotic chaperonin TRiC/CCT is essential for folding a diverse set of proteins, yet its interactome and functional roles in specialized neurons remain incompletely understood. To investigate TRiC-mediated folding in rod photoreceptors, we generated a transgenic mouse line expressing an epitope-tagged Tcp-1α subunit, enabling purification of intact TRiC complexes from retinal tissue. Mass spectrometry identified 226 TRiC-interacting proteins, including known TRiC substrates and co-chaperones as well as numerous novel candidates enriched in RNA processing, cytoskeletal organization, and cell-cycle regulation. Using a TRiC loss-of-function model in which expression of a short splice isoform of phosducin-like protein (PhLPs) competitively inhibits TRiC activity, we observed marked reductions in canonical TRiC substrates, including tubulins, transducin β subunits, and triosephosphate isomerase, as well as secondary alterations in proteins involved in cytoskeletal stability, membrane trafficking, energy metabolism, and phototransduction. Quantitative metabolomic profiling revealed that TRiC deficiency induces a metabolic “energy crisis” characterized by reduced glycolytic- and tricarboxylic acid cycle intermediates, acylcarnitines, ATP, NAD, and NADH, implicating widespread impairment of glucose utilization, mitochondrial bioenergetics, and fatty acid oxidation. Integrative proteomic–metabolomic analysis identified a small subset of proteins, including Rab10 and Anxa1, as potential drivers of these metabolic disruptions, with defective Rab10-dependent GLUT4 trafficking emerging as a plausible mechanism underlying impaired glucose uptake in TRiC-deficient rods. Finally, experiments using a perpetually unfolded Gβ_1_ mutant and Gγ_1_-knockout mice demonstrated that substrate overload sequesters TRiC and competitively displaces other clients, exacerbating proteostasis imbalance. Together, our study provides a comprehensive *in vivo* mapping of the TRiC interactome in mammalian rods, reveals a connection between TRiC-dependent proteostasis and energy metabolism in rods, and indicates a mechanism by which misfolded TRiC substrates exacerbate a proteostasis imbalance that ultimately results in neurodegeneration.

## Introduction

The visual function of vertebrates is mediated by rod and cone photoreceptors located in the retina, a specialized neural tissue at the back of the eye. Rod photoreceptors are responsible for peripheral and low-light night vision, while cone photoreceptors support high-acuity and color vision during the day. Both cell types rely on a specialized signaling cascade, phototransduction, which converts light into an electrical response. This process is initiated by the activation of rhodopsin, a G protein-coupled receptor, and its downstream effector, the heterotrimeric G protein transducin.

Phototransduction components are localized within stacked disc membranes of the photoreceptor outer segment—a modified primary cilium optimized for photon capture. The proper function and maintenance of this structure require continuous renewal and a high rate of protein synthesis and trafficking. Key phototransduction and structural proteins, including the transducin β subunit, actin, and tubulin, require specialized folding mechanisms to achieve their native conformations and avoid aggregation. In eukaryotic cells, this function is carried out by the cytosolic group II chaperonin known as TRiC (TCP-1 ring complex), also referred to as CCT (Chaperonin Containing TCP-1) [1].

Chaperonins are large, ∼1 MDa oligomeric protein complexes that facilitate ATP-dependent folding of non-native polypeptides. These complexes are conserved across all domains of life, with bacterial GroEL/GroES, archaeal thermosomes, and the eukaryotic TRiC/CCT forming three major families[2,3]. Structurally, chaperonins consist of two back-to-back stacked rings that create a central folding chamber[4,5]. Upon binding and hydrolysis of ATP, the rings undergo conformational changes that allow for encapsulation of the substrate protein, creating an isolated environment for productive folding[6].

TRiC is composed of eight distinct subunits (CCT1–CCT8), each with unique substrate specificities, which confer the complex a broad client repertoire. TRiC is essential for folding ∼10% of cytosolic proteins, particularly those with complex tertiary structures such as β-propellers and α/β-barrels[3,5,7]. Its known clients include tubulin, actin, G protein subunits, and key regulators of cell cycle, apoptosis, and transcription[8,9].

The importance of TRiC function in the nervous system, including the retina, is underscored by its disease associations. Mutations in TRiC subunits, such as *CCT2*, have been linked to Leber congenital amaurosis (LCA), a severe early-onset retinal dystrophy[10,11]. In addition, impaired TRiC function has been implicated in neurodegenerative diseases, including Alzheimer’s and Huntington’s diseases, where its expression is diminished in affected brain regions and during aging[12]. Other mutations in TRiC subunits lead to hereditary sensory and autonomic neuropathies (HSAN) and mutilating sensory neuropathies[13,14].

Proteomic profiling of the TRiC interactome have provided insight into its cellular functions and substrate diversity. These studies, performed in yeast[9], mammalian cell lines[3], and human brain tissue[15], have revealed a diverse array of TRiC client proteins involved in cytoskeletal organization, vesicular trafficking, proteostasis, and energy metabolism.

Here, we extended these findings by characterizing the TRiC interactome in the vertebrate retina. We employed an epitope-tagged TRiC complex (TRiC^et^) as a bait for affinity purification followed by mass spectrometry to identify TRiC-associated proteins in rod photoreceptors. We also analyzed global proteomic and metabolomic changes in a TRiC loss-of-function mouse model to identify affected pathways. Our results indicate that TRiC supports folding of proteins involved in diverse and essential processes, including cytoskeletal and membrane dynamics, mitochondrial and cytosolic energy metabolism, ATP-dependent chromatin remodeling, and RNA processing. Notably, we found that TRiC function can be competitively inhibited by a client protein carrying a folding-disruptive mutation, suggesting a potential pathogenic mechanism by which misfolded proteins interfere with chaperonin activity. Using converging *in vivo*–centered approaches, we identified protein interactors of TRiC in retinal photoreceptors and validated candidate substrates in their native cellular context. Our data provide new insights into the specific functions of TRiC, as well as its regulation and recycling in specialized mammalian neurons, offering a more accurate proteomic map of this essential chaperonin *in vivo*.

## Materials and Methods

### Animals models

The use of animal models in this study conforms to ARVO’s Statement for the Use of Animals in Ophthalmic and Vision Research. This study was approved by the West Virginia University Institutional Animal Care and Use Committee (protocol 1603001702).

#### TRiC^et^ mice

The coding sequence of *Mus musculus* Tcp-1α subunit of chaperonin was generated by Genscript. A FLAG-HA epitope tag flanked by GGG linker on both sides and an internal VKL linker, GGGDYKDDDDKVKLYPYDVPDYAGGG was inserted between the amino acid residues R^480^ and K^481^. This construct was used for transgene which contained a Kozak sequence at the 5’ end, a 4.4 kb mouse rhodopsin promoter[16], and a mouse protamine I polyadenylation sequence. The transgene was verified by sequencing and purified and injected into the pronuclei of zygotes from superovulated FVB females at West Virginia University. Transgene integration was confirmed by PCR genotyping of tail DNA using forward primer 5’-ACGACGATGACAAGGTCAAGCTCT and reverse primer 5’-ATCAATCCGAAGGATGGTGATT. Colonies were established by crossing transgenic heterozygotes with wild type partners of 129-E background (Charles River Laboratories).

PhLPs transgenic mice, rhodopsin P23H mice and Gγ_1_ knockout mice were previously described[17–20]. Each strain was back-crossed with 129-E mice for at least ten generations. The resulting isogenic strains were crossed with each other, and their offspring was ear-tagged and genotyped using PCR amplification of genomic DNA as described in the original papers. Mice were housed in adjacent cages, under standard diurnal cycle. Equal numbers of males and females were used for the analyses.

### TRiC^et^ pulldown and native PAGE

Frozen retinas (typically 20) or HEK293 cells pellet were homogenized in 0.8mL of 20 mM Hepes-HCl, pH 7.4, 100 mM KCl, 0.5 mM EDTA, 1% Triton X-100, and 10% glycerol by short ultrasonic pulses. The homogenate was cleared by centrifugation and the supernatant was incubated two times with 20µL of anti-FLAG agarose beads (A2220-1 ML; Sigma-Aldrich) for 30 minutes at room temperature. The beads were combined and washed one time with 0.5mL of the above buffer and two times with 0.5mL of 10 mM Hepes-HCl, pH 7.4. The captured proteins were eluted two times with 15µL of 10 mM Hepes-HCl, pH 7.4, 0.2mg/mL 3x FLAG peptide (F4799-4MG; Sigma) for 15 minutes. The eluates were combined and purified by Blue Native PAGE on a Novex 3-12% tris-glycine gel (BN2011BX10, Fisher) using a XCell SureLock Mini-Cell Electrophoresis System (EI0001; Fisher), at room temperature, according to manufacturer’s protocol. Gels were stained with Coomassie and a 900 kDa band was excised from the gel. The protein composition of the band was analyzed by LC MS/MS. In some cases, the band was incubated in 5% β-mercaptoethanol and 0. 5mL of 125 mM Tris/HCl, pH 6.8, 4% SDS, and 6M urea for 20 minutes at 95⁰C, placed in the well of a polyacrylamide gel, and analyzed by SDS-PAGE and Western blotting.

### Electroretinography (ERG)

Scotopic ERG responses were recorded and analyzed using either a Celeris (Diagnosis LLC) or an UTAS BigShot (LKC Technologies) rodent ERG system as previously described[21].

### Microscopy

#### Immunofluorescence microscopy

Confocal images of frozen retinal cross-sections were acquired as previously described[22].

#### Light microscopy and automated photoreceptor nuclei counting

Eyes were collected and sent to Excalibur Pathology, Inc (Norman, OK, USA) for histology processing. Total nuclei count within the outer nuclear layer (ONL) was determined in continuous ocular cross-sections stained with hematoxylin and eosin. The specimens were digitized using an Olympus VS120 slide scanner, and the resulting TIFF images were analyzed using a custom Cell Profiler pipeline, as previously described[21].

#### Negative staining transmission electron microscopy (TEM)

TRiC^et^ purified from HEK 293 cells was negatively stained with 1% uranyl acetate on carbon film coated copper grids. TEM images were obtained using a JEOL 1010 Transmission Electron Microscope equipped with an AMT Hamamatsu ORCA-HR Digital Camera. Particles were aligned and classified using SPIDER software image processing. Eyes were enucleated and immersed in fixative containing 2% paraformaldehyde, 2.5% glutaraldehyde, 100mM cacodylate buffer. After fixation, the eyes were transferred to a petri dish containing a drop of 7% sucrose buffer, and the cornea and lens were removed. The eyecups were then returned to the glass vial with the fixative for 48 hours with gentle rolling and then cut into smaller pieces. The pieces were washed five times for five minutes each with 0.1M cacodylate buffer at RT and then incubated with 2% osmium tetroxide in 0.1M cacodylate buffer for 1 hour on ice. After osmication, samples were washed with water and incubated with 1% aqueous uranyl acetate at 4°C overnight. The next day, after washes, samples were dehydrated in an ethanol series on ice. After dehydration, samples were washed with Propylene oxide 2 times for 15 minutes on a rotator at RT. Samples were then infiltrated with a 50:50 mixture of propylene oxide and resin for two hours at RT. Then, the solution was changed to 25:75 mixtures of propylene oxide and resin for overnight incubation at RT. The next day, the samples were transferred to fresh 100% resin and rotated for 2 hours at RT. Samples were then embedded in resin and cured in an oven at 60^0^ C overnight. Ultrathin sections (70 nm) were cut using a LEICA EM UC7 ultramicrotome, post-stained with Reynold’s lead citrate, and imaged the following day on a JEOL JEM-1400 transmission electron microscope.

### AAV vectors design, production, and subretinal injections

The coding sequences of mouse phosducin-like protein (*Pdcl*) or Gβ_1_ (*Gnb1*) were synthesized by Genscript and inserted into pcDNA3.1+. PhLPs-6x-His consisted of the cDNA sequence encoding Pdcl with DNA bases corresponding to amino acids 1-83 removed and a 6x-His tag added to the C-terminus. ^Δ130-136^ PhLPs-6x-His contains additional deletion of amino acids 130-136. Myc-Gβ_1_ consisted of the cDNA sequence encoding *Gnb1* with a myc-tag added to the N-terminus. Myc-Gβ1AA contained two alanine insertions after Leu7 and Ala11. Each construct was placed under the mouse opsin promoter (mOP), which limits the expression to rod photoreceptors, and packaged into AAV-PHP.eB capsid, as previously described[21]. The AAV production was conducted at the Viral Core of West Virginia University. The subretinal AAV injection was performed using 1-2-month-old 129-E mice, as previously described[21].

### Tissue culture

Plasmids for the expression of human tcp-1α^et^, PhLPs-6x-His, ^Δ130-136^ PhLPs-6x-His, myc-Gβ_1_, myc-Gβ_1_^AA^ and HA-Gγ_1_ in mammalian cell culture were made by GenScript. HEK 293 cells were maintained in DMEM/F-12 medium supplemented with 10% FBS and 1% penicillin-streptomycin in an incubator at 37⁰C and 5% CO_2_. For transient protein expression, cells were plated in six-well plates and grown to 80% confluency. The following day, cells were transfected with 2.5 μg of plasmid DNA using polyethylenimine (PEI) at a 1 (plasmid DNA): 2.5 (PEI) ratio. Empty pcDNA3.1+ vector was used as a control. Cells were collected 48h after transfection, frozen on dry ice, and stored at -80⁰C.

### Antibodies

Proteins were detected using the following antibodies: rat anti-HA (Roche, 11867423001), rabbit anti-FLAG (Proteintech, 20543-1-AP), mouse anti-tcp-1β (Santa Cruz Biotechnology, sc-374152), rat anti-tcp-1α (Santa Cruz Biotechnology, sc-53454) mouse anti-6x-His tag (MA1-21315, Invitrogen).

### Mass spectrometry

#### DDA Mass spectrometry at Duke University

The 900 kDa band was de-stained, and subjected to in-gel tryptic digestion[23]. The resulting peptides were extracted from the gel and vacuum dried. Peptide mixes were dissolved in 12μL of 1/2/97% (by volume) of the trifluoroacetic acid/acetonitrile/water solution, and 3μL was injected into a 5μm, 180μm×20mm Symmetry C18 trap column (Waters) in 1% acetonitrile in water for 3 min at 5μL/min. The analytical separation was next performed on an HSS T3 1.8μm, 75 μm×200 mm column (Waters) over 60 min at a flow rate of 0.3 μL/min at 55°C using nanoAcquity UPLC system (Waters Inc.). The 5-30% mobile phase B gradient was used, where phase A was 0.1% formic acid in water and phase B 0.1% formic acid in acetonitrile. Peptides separated by LC were introduced into the Q Exactive HF Orbitrap mass spectrometer (Thermo Fisher Scientific) using positive electrospray ionization at 2000 V and capillary temperature of 275°C. Data collection was performed in the data-dependent acquisition (DDA) mode with 120,000 resolution (at m/z 200) for MS1 precursor measurements. The MS1 analysis utilized a scan from 375-1500 m/z with a target AGC value of 1.0e6 ions, the RF lens set at 30%, and a maximum injection time of 50ms. Advanced peak detection and internal calibration (EIC) were enabled during data acquisition. Peptides were selected for MS/MS using charge state filtering (2-5), monoisotopic peak detection and a dynamic exclusion time of 20 sec with a mass tolerance of 10 ppm. MS/MS was performed using HCD with a collision energy of 30±5% with detection in the ion trap using a rapid scanning rate, AGC target value of 5.0e4 ions, maximum injection time of 150 ms, and ion injection for all available parallelizable time enabled.

Next, label-free relative protein quantification was performed with raw mass spectral data files (.raw) were imported into Progenesis QI for Proteomics 4.2 software (Nonlinear Dynamics) for duplicate runs alignment of each preparation and peak area calculations. Peptides were identified using Mascot version 2.5.1 (Matrix Science) for searching the UniProt reviewed mouse database. Mascot search parameters were: 10 ppm mass tolerance for precursor ions; 0.025 Da for fragment-ion mass tolerance; one missed cleavage by trypsin; fixed modification was carbamidomethylation of cysteine; variable modification was oxidized methionine. To account for variations in experimental conditions and amounts of protein material in individual LC-MS/MS runs, the integrated peak area for each identified peptide was corrected using the factors calculated by automatic Progenesis algorithm utilizing the total intensities for all peaks in each run. Values representing protein amounts were calculated based on a sum of ion intensities for all identified constituent non-conflicting peptides. To determine protein abundances, two duplicate runs for each sample were averaged and ratio between the compared samples was calculated and log2 normalized in R.4.1.2. Proteins associated with TRiC were defined as significantly enriched if they were identified by at least two unique peptides and exhibited an absolute log2-fold change ≥ 1 with an ANOVA-adjusted *p-value < 0.05*.

#### DIA mass spectrometry at IDeA National Resource for Quantitative Proteomics, University of Arkansas

Proteins from in the 900 kDa band were reduced, alkylated, and digested in-gel with sequencing grade modified porcine trypsin (Promega). Tryptic peptides were then separated by reverse phase XSelect CSH C18 2.5 um resin (Waters) on an in-line 150 x 0.075 mm column using an UltiMate 3000 RSLCnano system (Thermo). Peptides were eluted using a 60 min gradient from 98:2 to 65:35 buffer A:B ratio (buffer A = 0.1% formic acid, 0.5% acetonitrile; buffer B = 0.1% formic acid, 99.9% acetonitrile). Eluted peptides were ionized by electrospray (2.2 kV) followed by mass spectrometric analysis on an Orbitrap Exploris 480 mass spectrometer (Thermo). To assemble a chromatogram library, six gas-phase fractions were acquired on the Orbitrap Exploris with 4 m/z DIA spectra (4 m/z precursor isolation windows at 30,000 resolution, normalized AGC target 100%, maximum inject time 66 ms) using a staggered window pattern from narrow mass ranges using optimized window placements. Precursor spectra were acquired after each DIA duty cycle, spanning the m/z range of the gas-phase fraction (i.e. 496-602 m/z, 60,000 resolutions, normalized AGC target 100%, maximum injection time 50 ms). For wide-window acquisitions, the Orbitrap Exploris was configured to acquire a precursor scan (385-1015 m/z, 60,000 resolution, normalized AGC target 100%, maximum injection time 50 ms) followed by 50x 12 m/z DIA spectra (12 m/z precursor isolation windows at 15,000 resolution, normalized AGC target 100%, maximum injection time 33 ms) using a staggered window pattern with optimized window placements. Precursor spectra were acquired after each DIA duty cycle.

Following data acquisition, data were searched using a corrected library against the UniProt *Mus musculus* database (June 2021) and a quantitative analysis was performed to obtain a comprehensive proteomic profile. Proteins were identified and quantified using EncyclopeDIA[24] and visualized with Scaffold DIA using 1% false discovery thresholds at both the protein and peptide level. Protein MS2 exclusive intensity values were assessed for quality using ProteiNorm[25]. The data was normalized using cyclic loess[26] and analyzed using R studio v1.2.0 with the package proteoDA to perform statistical analysis using Linear Models (limma) with empirical Bayes (eBayes) smoothing to the standard errors [26]. Proteins with an FDR < 0.05 and absolute fold change ≥ 1 were considered significant.

#### TMT Mass spectrometry Orbitrap Eclipse

For TMT-MS, total protein from each sample was reduced, alkylated, and purified by chloroform/methanol extraction prior to digestion with sequencing grade trypsin (Promega). The resulting peptides were labeled using a tandem mass tag 11-plex isobaric label reagent set (Thermo) and combined into one multiplex sample group. Labeled peptides were separated into 46 fractions on a 100 x 1.0 mm Acquity BEH C18 column (Waters) using an UltiMate 3000 UHPLC system (Thermo) with a 50 min gradient from 99:1 to 60:40 buffer A:B ratio under basic pH conditions, then consolidated into 18 super-fractions (Buffer A = 10 mM ammonium hydroxide, 0.5% acetonitrile; Buffer B = 10 mM ammonium hydroxide, 99.9% acetonitrile; both buffers adjusted to pH 10 for offline separation.

Each super-fraction was then further separated by reverse phase XSelect CSH C18 2.5 um resin (Waters) on an in-line 150 x 0.075 mm column using an UltiMate 3000 RSLCnano system (Thermo). Peptides were eluted using a 75 min gradient from 98:2 to 60:40 buffer A:B ratio (buffer A = 0.1% formic acid, 0.5% acetonitrile; buffer B = 0.1% formic acid, 99.9% acetonitrile). Eluted peptides were ionized by electrospray (2.2 kV) followed by mass spectrometric analysis on an Orbitrap Eclipse Tribrid mass spectrometer (Thermo) using multi-notch MS3 parameters. MS data were acquired using the FTMS analyzer in top-speed profile mode at a resolution of 120,000 over a range of 375 to 1500 m/z. Following CID activation with normalized collision energy of 31.0, MS/MS data were acquired using the ion trap analyzer in centroid mode and normal mass range. Using synchronous precursor selection, up to 10 MS/MS precursors were selected for HCD activation with normalized collision energy of 55.0, followed by acquisition of MS3 reporter ion data using the FTMS analyzer in profile mode at a resolution of 50,000 over a range of 100-500 m/z.

#### TMT Mass Spectrometry Data Analysis

Proteins were identified and MS3 reporter ions quantified using MaxQuant (Max Planck Institute) against the UniprotKB *Mus musculus* ( 2023) database with a parent ion tolerance of 3 ppm, a fragment ion tolerance of 0.5 Da, and a reporter ion tolerance of 0.003 Da. Scaffold Q+S (Proteome Software) was used to verify MS/MS based peptide and protein identifications were accepted if they could be established with less than 1.0% false discovery and contained at least 2 identified peptides; protein probabilities were assigned by the Protein Prophet algorithm[27] and to perform reporter ion-based statistical analysis. Protein TMT MS3 reporter ion intensity values are assessed for quality and normalized using ProteiNorm[25]. The data was normalized using Cyclic Loess[26] and differential expression analysis was performed as mentioned above in R studio v12.0 with the package proteoDA. Differentially expressed proteins were considered significantly altered if they exhibited an adjusted p-value < 0.05 and an absolute fold change ≥ 1.5; to avoid bias, any proteins annotated as “Cct” or “tcp1” (known TRiC complex components) were excluded from the analysis.

#### Functional enrichment analysis

Relevant biological processes and interaction networks associated with TRiC interactors were compiled by pooling proteins identified in both DDA and DIA experiments, then subjecting this set to enrichment analysis of Gene Ontology Biological Process, Molecular Function and Cellular Component terms, as well as KEGG pathways, using Metascape[28] as a meta-analysis platform. Biochemical characteristics of TRiC interactors such as WD40 repeats, β-propeller folds and other domain architectures were retrieved by custom R (v4.2.1) scripts querying the InterPro database. For proteins differentially expressed in the TMT-MS workflow, Gene Set Enrichment Analysis (GSEA) was performed with ClusterProfiler[29] v4.0 to identify enriched GO Biological Process and Molecular Function categories. Data visualizations, volcano plots illustrating differential protein interactors and heatmaps were generated in R (v4.1.2) within RStudio v1.2.0 using the EnhancedVolcano and pheatmap packages.

#### Metabolomic analysis

Targeted metabolomics was performed as previously described [30,31]. Briefly, flash frozen mouse retinas were homogenized in 80% cold methanol to extract metabolites. Supernatants containing aqueous metabolites were dried and analyzed by liquid chromatography–mass spectrometry (LC MS). An ACQUITY UPLC A BEH Amide analytic column (2.1 × 50 mm, 1.7 μm, Waters) in a Shimadzu LC Nexera X2 UHPLC was used for chromatographic separation and metabolites were detected with a QTRAP 5500 LC MS (AB Siex). The extracted multiple reaction monitoring (MRM) peaks were integrated using MultiQuant 3.0.2 software (AB Sciex). The pellets of the extraction were analyzed for protein concentration for sample normalization.

#### Metabolomic Data Analysis

Each treated PhLPs and Rho P23H group was compared with wild type 128E group, using Metaboanalyst 6.0 for statistical data analysis.

#### Multi-omics integration via DIABLO

Proteomics and metabolomics data were first pre-processed independently in R.4.2.1. Proteomic intensities (log₂-cyclic-loess normalized) were filtered to retain proteins with |log₂ fold-change| > 0.59 and P < 0.05 between PhLPs and wild type 129E controls. Metabolomic intensities (log₂-transformed) were quantile-normalized without imputation, and metabolites whose PhLPs vs 129E and P23H vs 129E fold-changes pointed in opposite directions (regardless of magnitude) were selected. Both data blocks were subset to the same ten samples (five PhLPs, five 129E), and their row names aligned. Using the mixOmics R package, a fully-connected two-block design matrix (off-diagonal = 1, diagonal = 0) was defined. We then performed a 5-fold cross-validation with the function (tune.block.splsda) over candidate keepX grids (proteins: 5, 10, 15, 20; metabolites: 10, 20, 30, 40) to identify the optimal number of features per block on two components. The final DIABLO model function (block.splsda) was fit with the chosen keepX and design, and model performance assessed via classification error rates. Top multi-omics signatures were visualized by loading barplots (plotLoadings), clustered image maps (cimDiablo) to reveal strongly correlated protein–metabolite modules.

## Results and Discussion

### Epitope-tagging of TRiC in rod photoreceptors

To enable affinity purification of the chaperonin complex, an epitope tag was inserted into the coding sequence of the tcp-1α subunit at a protruding equatorial loop (Fig 1A). The epitope-tagged tcp-1α transgene, driven by a rod photoreceptor-specific promoter, was expressed in mice. Transgene expression was confirmed by Western blotting, either through anti-FLAG pulldown or direct probing of retinal extracts (Fig 1BC). The combined fluorescence of transgenic and endogenous tcp-1α bands in transgene-positive retinas matched the fluorescence of the tcp-1α band in wild-type retinas, indicating no significant overexpression in transgenic mice (Fig 1C).

**Fig 1.**
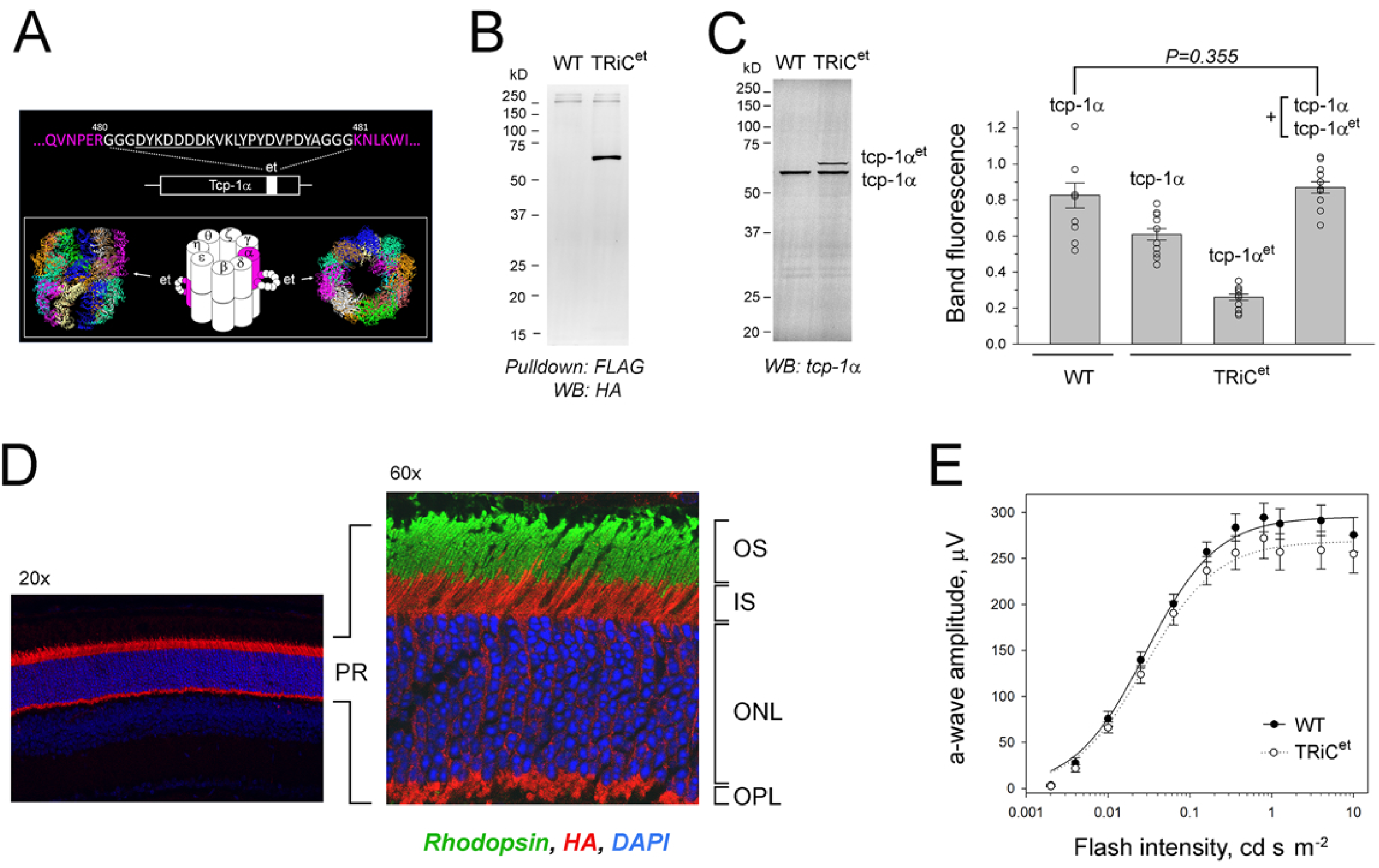
Epitope-tagging of chaperonin in rod photoreceptors. ***A***. DYKDDDK (FLAG) and LYPYDVPDYA (HA) sequences inserted into tcp-1α (magenta) transgene expressed in rods of TRiC^et^ mice. Resulting epitope tags (et), illustrated on a TRiC^et^ cartoon, protruded from the loop indicated with arrow on side- and top TRiC view with color-coded subunits (PDB ID 6QB8). ***B***. TRiC^et^ detected by anti-FLAG pulldown and Western blotting in the retinas of wild-type (WT) and TRiC^et^ mice. ***C***. The level of the endogenous and epitope-tagged tcp-1α quantified by Western blotting (SEM, n=11). Significance was determined by Mann-White rank sum test. ***D***. TRiC^et^ visualized in retinal cross-section by immunofluorescence confocal microscopy with rat anti-HA (orange); PR-photoreceptors, OS-outer segments, IS-inner segments, ONL -outer nuclear layer, OPL-outer plexiform layer. Staining with mouse anti-rhodopsin (green) and DAPI (blue) are included on 60x panel as markers of rod outer segments and nuclei. No anti-HA staining was detected in wild-type retinas (not shown). ***E***. Visual responses of dark-adapted rods were recorded using a Celeris (Diagnosis LLC) rodent ERG system. The amplitude of elicited ERG a-wave is shown as a function of flash intensity (SEM, n= 8). Responses of wild-type (black symbols) and TRiC^et^ (white symbols) mice were compared at two months of age. Significance was determined using t-test. Each data set was fitted with a simple rectangular hyperbola with two parameters using SigmaPlot 13 software.

The transgene was uniformly expressed in rod photoreceptors and localized to all subcellular compartments except the outer segment (Fig 1D). Importantly, its expression had no adverse effect on rod visual function (Fig 1E). To confirm that the epitope-tagged tcp-1α assembled into TRiC, it was pulled down using anti-FLAG agarose, eluted under non-denaturing conditions, and analyzed by native gel electrophoresis (Fig 2A). Additionally, the tcp-1α construct used for transgene production was transiently expressed in HEK293 cells. Pulldowns from both the cell culture and retinal extracts ran as a ∼900 kDa band, closely resembling TRiC isolated from the retinas of PhLPs mice. Negative-stain TEM further confirmed that the captured complex had the characteristic ring-shaped structure, similar to the endogenous TRiC captured by its co-factor PhLP (Fig 2BC). Based on these results, the generated transgenic strain was named TRiC^et^.

**Fig 2.**
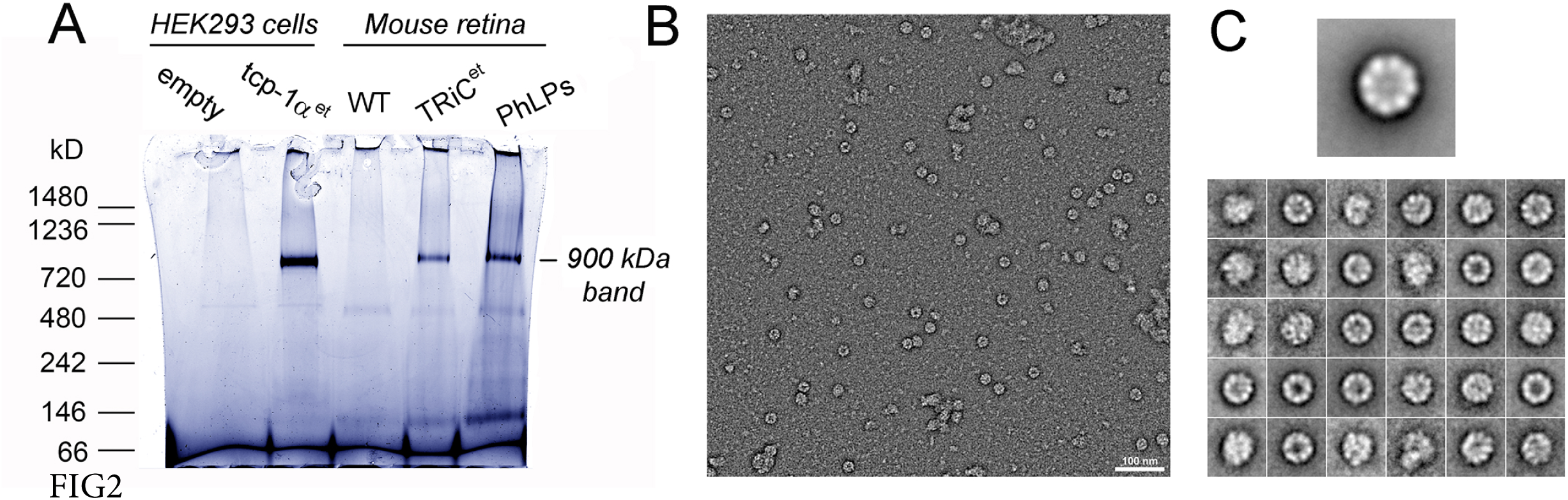
Chaperonin complex runs on a Blue Native gel as the 900kDa band. ***A***. Anti-FLAG pulldown of TRiC^et^ under non-denaturing conditions separated on Blue Native Gel and stained with Coomassie. HEK293 cells were transfected with empty vector or tcp-1α^et^ vector; retinas were collected from wild type 129E mice, TRiC^et^ and PhLPs mice at 21 days of age. ***B***. Negative stain TEM of anti-FLAG pulldown from HEK293 cells transfected with PhLP-FLAG showing ring-shaped particles that were absent in the pulldown from HEK293 cells transfected with empty vector (not shown). ***C***. Global average image of 513 particle images aligned showing the most common top view TRiC orientation with eight subunits visible (top), and the same data classified into 30 classes (bottom).

### TRiC loss-of-function model

Phosducin-like protein (PhLP, gene: Pdcl1) is a co-chaperone that facilitates the folding of Gβ and binds chaperonin with high affinity[32]. Previously, we found that the short splice isoform of PhLP, PhLPs, which lacks the Gβ-binding domain, induces rod outer segment breakdown and cell death[17,18,33]. We proposed that PhLPs causes this phenotype by inhibiting the folding of nascent proteins by chaperonin.

To further support this mechanism, we deleted amino acids 130–136 of PhLPs, a region implicated in TRiC binding (Fig 3A)[34]. This mutant, termed ^Δ130-136^PhLPs-6x-His, and the control PhLPs-6x-His were co-expressed with epitope-tagged Tcp-1α in HEK293 cells, followed by TRiC^et^ pulldown assays. Western blot analysis of the pulldowns, both directly and after isolating the 900 kDa TRiC^et^ band by native gel electrophoresis (Fig 3B), showed that PhLPs-6x-His co-precipitated with TRiC^et^ more efficiently than ^Δ130-136^PhLPs-6x-His. Importantly, the expression levels of the transfected proteins and endogenous chaperonin were comparable. These results confirm that the Δ130-136 deletion reduces PhLPs binding to chaperonin.

**Fig 3.**
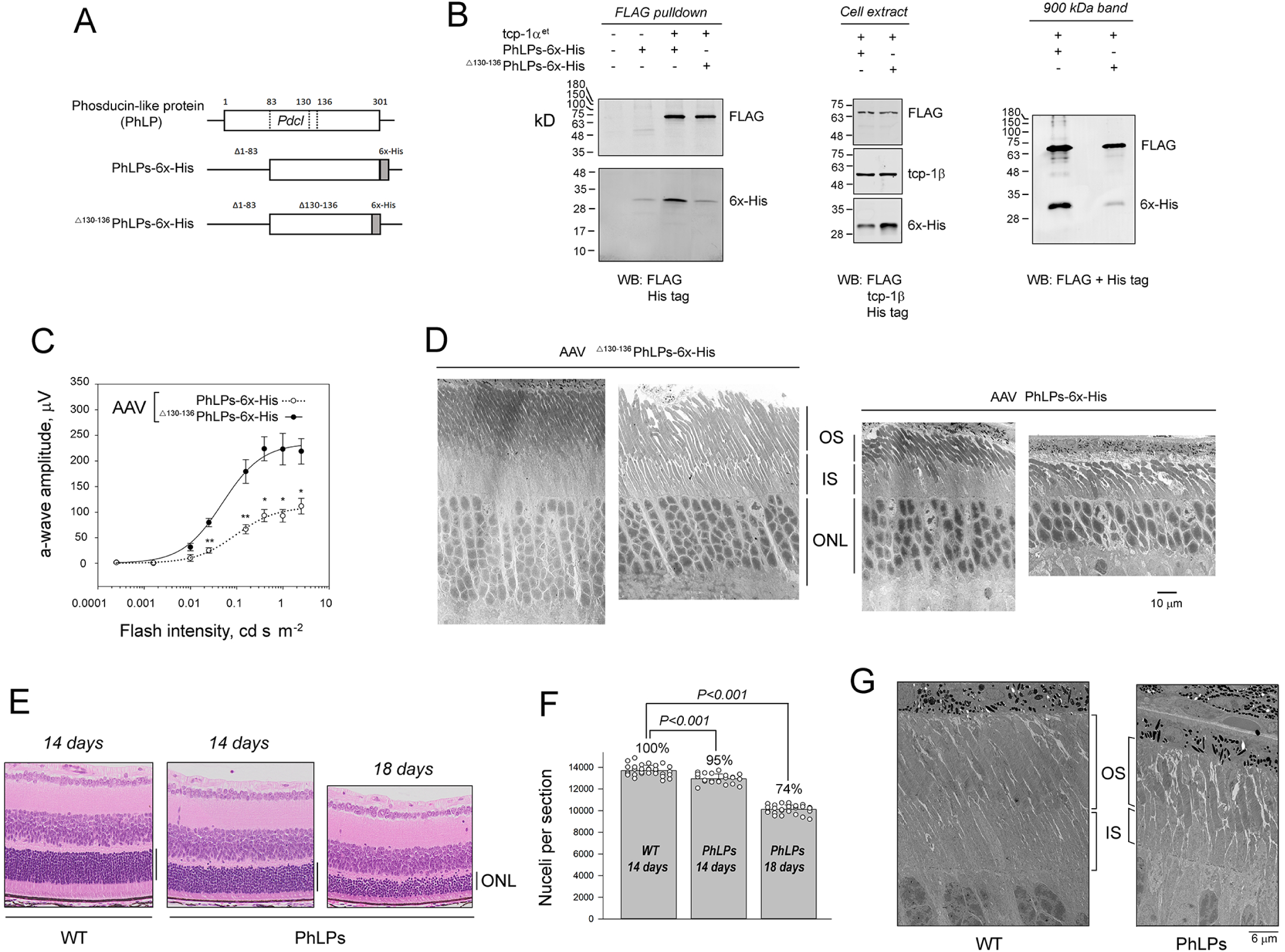
PhLPs binding to TRiC leads to rod photoreceptors death. ***A***. Phosducin-like protein (gene: *Pdcl*; protein: PhLP) and its epitope-tagged short splice isoform PhLPs-6x-His and ^Δ130-136^PhLPs-6x-His mutant. ***B***. Western blot analysis of HEK293 cells co-transfected with tcp-1α^et^ and PhLPs-6x-His or ^Δ130-136^PhLPs-6x-His: anti-FLAG pulldown (left), whole cell extract (center), the 900kDa band from Blue Native Gel of anti-FLAG pulldown (right). ***C***. Comparison of visual responses of mice that received subretinal injection of AAV-PhLPs-6x-His (white circles) and AAV -^Δ130-136^PhLPs-6x-His (black circles) 2-months post-injection. Electroretinographic responses were recorded using a UTAS BigShot (LKC Technologies) rodent ERG system. The amplitude of elicited ERG a-wave is plotted as a function of flash intensity (SEM, n= 4). Each dataset was fitted with a simple rectangular hyperbola with two parameters. The significance was determined by t-test and Mann-White rank sum test, with P value <0.05 (*) and <0.01 (**). ***D***. Ultrastructure of rod photoreceptors of mice in *C* analyzed by TEM: outer segments (OS), inner segments (IS), outer nuclear layer (ONL). E. Ocular cross-section stained with hematoxylin and eosin of wild type 129E mice at 14 days of age, and PhLPs mice at 14 and 18 days of age. ONL: outer nuclear layer containing stacked photoreceptors’ nuclei. ***F***. ONL nuclei counts in the entire retinal cross-sections measured by ONLyzer: bars are mean value with SD, P-value determined by t-test, n=31(WT), 22 (PhLPs, 14 days-old), 28 (PhLPs, 18 days-old). ***G***. Ultrastructure of rod photoreceptors at 14 days of age analyzed by TEM: outer segments (OS), inner segments (IS).

Next, we compared the *in vivo* effects of PhLPs-6x-His and ^Δ130-136^PhLPs-6x-His using an AAV delivery system. The amplitude of the a-wave recorded in dark-adapted mice via electroretinography served as a measure of total number of rods in the retina and phototransduction efficiency in their outer segments (Fig 3C). Rod ultrastructure was analyzed by TEM (Fig. 3D). Mice treated with AAV: ^Δ130-136^PhLPs-6x-His exhibited significantly larger a-wave amplitudes, retained more rod nuclei in the outer nuclear layer (ONL), and had longer rod outer segments than those treated with AAV: PhLPs-6x-His. These findings suggest that the toxicity of PhLPs is attenuated when TRiC binding is disrupted, supporting the hypothesis that TRiC is a primary target of PhLPs in rods.

To determine the onset of rod photoreceptor degeneration, we analyzed transgenic PhLPs mice, which begin expressing PhLPs-FLAG in rods at postnatal day 8. The total number of photoreceptor nuclei within the ONL was quantified using hematoxylin and eosin-stained paraffin-embedded retinal cross-sections (Fig 3EF). PhLPs-expressing mice and wild-type controls of the same genetic background were compared at postnatal days 14 and 18. At day 14, 95% of PhLPs-expressing photoreceptors were still present, but this percentage declined to 74% by day 18. Additionally, PhLPs mice displayed shortened rod outer and inner segments as early as day 14 (Fig 3G).

These results further support the role of PhLPs in disrupting chaperonin-mediated protein folding, leading to rod photoreceptor degeneration. Importantly, our findings suggest that postnatal day 14 and possibly 18 represent key time points when TRiC function is already fully suppressed by PhLPs, yet rods have not undergone widespread cell death. This provides an opportunity to analyze proteome changes associated with TRiC loss-of-function before major degeneration occurs in conjunction with the analysis of the retina specific TRiC^et^ interactome.

### Identification of Proteins Interacting with TRiC^et^ in Mouse Retina

To identify proteins interacting with TRiC^et^ in rod photoreceptors, this protein complex was captured using an anti-FLAG pulldown, separated as a 900 kDa band on native gel electrophoresis, and analyzed by LC-MS/MS. Pulldowns from age-matched wild-type retinas processed under identical conditions served as the baseline. The abundance of identified proteins was expressed as the fold change in TRiC^et^ versus wild-type samples. To account for potential instrumentation bias, two independent datasets from 2 orthogonal approaches, DDA and DIA were generated at the proteomic facilities of Duke University and the IDeA Proteomic Facility at the University of Arkansas respectively.

In both datasets, the tcp-1 subunits of chaperonin were identified with high confidence, reinforcing that the 900 kDa band represented an intact chaperonin complex. Additionally, the Duke and IDeA datasets contained 131 and 109 proteins, respectively, that were enriched by more than twofold (Sup. Table I). The overlap between these datasets was limited, with only 14 shared proteins, six of which were known TRiC substrates (Gnb1, Gnb2, Gnb5, Hdac1) or TRiC co-chaperones (Pdcl, Pdcl3) (Fig 4AB). However, each dataset included additional known TRiC interactors that were not detected by any specific data filter. Consequently, both datasets were merged into a single database containing 226 targets for subsequent analyses (Sup. Table II).

**Fig. 4.**
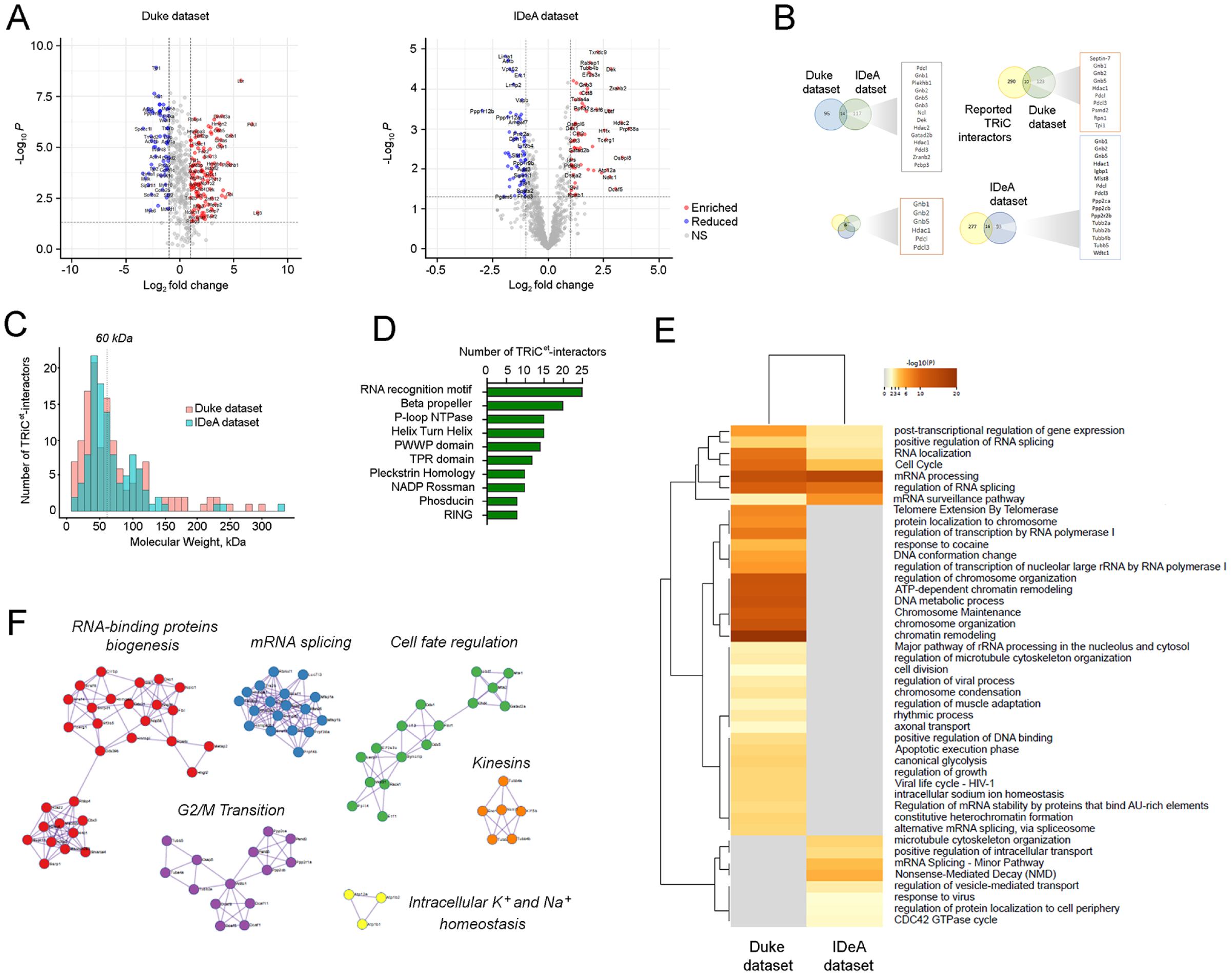
Analysis of TRiC^et^ interactome in mouse retina. ***A***. Volcano plots displaying differential proteins of the 900 kDa band in preparations from TRiC^et^ and wild type retinas. The x-axis represents the log₂ fold change (log₂FC) and the y-axis shows (–log₁₀(p-value), with p < 0.05 and |log₂FC| > 1 inclusion criteria for all identified proteins. The chaperonin subunits were removed from the plot. Shown datasets were independently obtained at the proteomic facilities of Duke University and the IDeA Proteomic Facility at the University of Arkansas. ***B***. Venn diagrams illustrating the overlap between the Duke and IDeA datasets (top) and previously published TRiC interactors (bottom) with gene names tabulated. ***C***. Distribution of molecular weights in across both mass spec datasets of TRiC^et^ interactors. The 60 kDa line indicates the theoretical size limit of the TRiC folding chamber. ***D***. Domain architecture frequency analysis in the combined dataset of TRiC^et^ interactors using InterPro/Pfam searches. ***E***. Heatmap illustrating the enrichment comparison of both KEGG and Reactome pathways in the Duke and IDeA datasets as determined by Metascape enrichment meta-analysis. The pathway significance is encoded using a gradient based on –log₁₀(p-value), highlighting statistically significant pathways associated with TRiC interactors gray rows indicate no enrichment found. ***F***. Protein–protein interaction networks identified in the combined dataset of TRiC^et^ interactors and the related biological process.

The structural and physicochemical properties of these 226 TRiC^et^-interacting proteins were examined, revealing that approximately 60% (130 out of 226) had molecular weights below 60 kDa (Fig. 4C), aligning with the size constraints of TRiC’s folding chamber. The most common structural features included RNA recognition motifs and beta-propeller folds (Fig 4D). Many identified proteins were found to participate in mRNA processing and regulation of cell cycle (Fig 4EF). We concluded that, in addition to *bona fide* TRiC substrates encapsulated in the folding cavity, our dataset includes proteins that regulate TRiC activity. To determine which proteins are directly folded by TRiC, we analyzed changes in steady-state protein levels in the retina in response to PhLPs transgene expression, which suppresses TRiC-mediated protein folding.

### Changes in Protein Levels in the Retina of PhLPs Mice

Retinas of PhLPs mice were collected at postnatal days P14 and P18, and steady-state protein levels were compared to those in age-matched wild-type mice using isobaric labeling with tandem mass tags TMT-MS (Sup. Table III). At P14, 24 proteins were significantly reduced in PhLPs retinas, including 11 TRiC^et^ interactors (Fig 5AB). The largest group consisted of tubulin family proteins (Tubb5, Tubb4a, Tubb2a, Tubb2b, Tubb4b, and Tuba4a). Given that alpha- and beta-tubulins are well-established TRiC substrates, this finding supports the notion that PhLPs inhibit TRiC-mediated tubulin folding, leading to its accelerated proteolytic degradation. This observation provides a compelling explanation for the severe underdevelopment of the rod outer segment in PhLPs mice. The rod outer segment, a microtubule-based compartment, undergoes rapid growth at P14, and its development is arrested when TRiC-mediated tubulin folding is disrupted.

**Fig 5.**
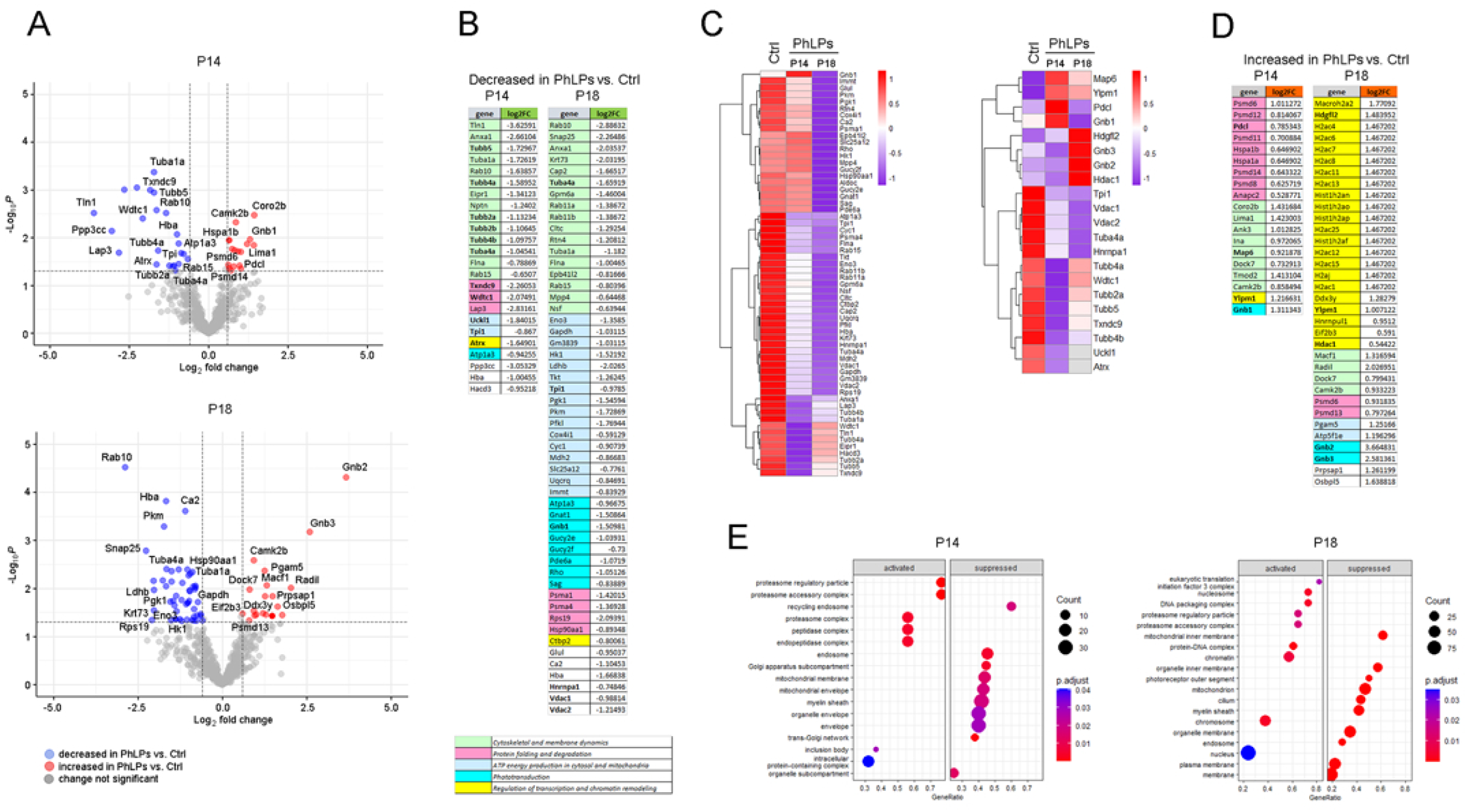
Changes of retina proteome caused by the PhLPs transgene. ***A***. Volcano plots showing differential protein levels in the retina of PhLPs mice vs. wild type controls 14- and 18-days after birth (P14 and P18, respectively). The x-axis represents the log₂ fold change (log₂FC) and the y-axis shows (–log₁₀(p-value), with p < 0.05 and |logFC| > 1.5 inclusion criteria. The data were collected at the IDeA Proteomic Facility at the University of Arkansas. ***B***. Proteins decreased in the retina of PhLPs mice grouped and color-coded based on their biological function. The identified TRiC^et^ interactors (Table II) are in bold font. ***C***. Heatmap of differentially expressed proteins (p < 0.05) across the entire PhLPs dataset (left) and the identified TRiC^et^ interactors (right), with z-scores indicating the relative expression (red: upregulated; blue: downregulated). ***D***. Proteins increased in the retina of PhLPs mice grouped and color-coded based on their biological function. The identified TRiC^et^ interactors are in bold font. ***E***. Enrichment of GO biological processes in the PhLPs dataset assessed by GSEA using ClusterProfiler. Each dot represents a GO term, with its position on the x-axis reflecting the normalized enrichment score (NES), positive NES values denote processes up-regulated/activated in the differential protein set, while negative NES values denote down-regulated/suppressed processes. Dot size corresponds to the number of proteins contributing to that term, and color shading indicates statistical significance (adjusted p< 0.05).

Another confirmed TRiC^et^ interactor that was reduced in PhLPs retinas was thioredoxin domain-containing protein 9 (Txndc9), a homolog of phosducin-like protein. Txndc9 has been proposed to negatively regulate TRiC ATPase activity and the folding of nascent tubulin and actin[35], which may explain its reduced levels in rods experiencing tubulin deficiency. Three additional TRiC^et^-interacting proteins, WD and tetratricopeptide repeat-containing protein 1 (Wdtc1), uridine-cytidine kinase-like 1 (Uckl1), and triosephosphate isomerase (Tpi1, were cytosolic proteins under 80 kDa, suggesting they may also serve as TRiC substrates. Additionally, Wdtc1, which has been proposed to act as a substrate receptor for CRL4 E3 ligase complex mediating ubiquitin-dependent protein degradation[36], may play a role in regulating TRiC turnover. A novel TRiC partner identified in this study was ATRX, a 279 kDa transcriptional regulator involved in chromatin remodeling and transcriptional control.

Interestingly, several non-TRiC interactors that were notably reduced at P14 clustered around Ca²⁺ signaling and homeostasis. These included neuroplastin (Nptn), a chaperone of plasma membrane Ca²⁺ ATPase; Ca²⁺-dependent protein phosphatase (Ppp3cc); the metallopeptidase Lap3; and annexin A1 (Anxa1), a regulator of phagosome-actin cytoskeleton interactions. Another downregulated group consisted of proteins involved in intracellular membrane trafficking (e.g., Rab10, Eipr1, Snap25, Rab15) and actin cytoskeleton regulation (Tln1, Cdc42, Flna). Additionally, we observed a reduction in Hacd3, an enzyme catalyzing long-chain fatty acid elongation in the endoplasmic reticulum, and Atp1a3, a Na⁺/K⁺-ATPase essential for maintaining membrane potential in visual signaling.

By P18, the number of proteins reduced in PhLPs retinas increased to 53 (Fig 5AB). However, only eight of these proteins were confirmed TRiC^et^ interactors, including tubulin (Tuba4a) and triosephosphate isomerase (Tpi), whose reductions were also observed at P14. The observed reduction of rod-specific transducin beta subunit (Gnb1) supports previous findings that Gnb1 is a major TRiC substrate affected by PhLPs[17]. Additionally, heterogeneous nuclear ribonucleoprotein A1 (Hnrnpa1) displayed characteristics of a potential TRiC substrate. However, this was deemed unlikely for two membrane proteins (Vdac1, Vdac2), as TRiC primarily folds soluble substrates.

Interestingly, the destabilization of two potential TRiC substrates, Uckl1 (involved in the pyrimidine salvage pathway) and Tpi1 (a key enzyme in glycolysis), triggered widespread effects on protein networks related to glucose metabolism and energy production. As a result, several non-TRiC-interacting proteins (e.g., Ldhb, Pfkl, Gapdh, Pkm, Pgk1, Hk1, Eno3, Tkt, Aldoc, Cox4i1, Cyc1, Mdh2, Slc25a12, Uqcrq, and Immt) were decreased at P18 (Fig 5B). Similarly, disruption of tubulin folding led to secondary effects on cytoskeletal proteins (e.g., Cap2, Krt73, Epb41l2, Flna), membrane trafficking proteins (e.g., Rab10, Rab11b, Rab11a, Rab15, Snap25, Anxa1), and phototransduction proteins involved in visual signaling in the outer segment (e.g., Gnat1, Gucy2e, Gucy2f, Pde6a, Rho, Sag, Atp1a3). However, none of these proteins appeared to directly interact with TRiC^et^. These findings suggest that the observed changes in the retinal proteome at P18 increasingly reflect secondary effects stemming from the destabilization of TRiC substrates at P14.

In addition, we observed adaptive responses resulting from the suppression of protein folding by TRiC. This category included proteins whose levels in the retina increased in response to the PhLPs transgene (Fig 5CD). For example, at P14, the levels of 19 proteins were significantly elevated in the retinas of PhLPs mice. Among these, four were confirmed TRiC interactors, including protein involved in cytoskeletal maintenance (Map6), a DNA-binding regulator of telomerase (Ylpm1), the rod-specific transducin beta subunit (Gnb1), and phosducin-like protein (Pdcl).

When categorizing all upregulated proteins based on biological function, they clustered into two major groups: proteins involved in maintaining proteostasis (e.g., Psmc5, Psmd6, Psmd8, Psmd11, Psmd12, Psmd14, Hspa1a, and Hspa1b) and regulators of cytoskeletal dynamics (e.g., Coro2b, Lima1, Tmod2, Ina, Map6, Dock7, and Ank3). This pattern suggests that the disruption of proteostasis and cytoskeletal dynamics may be the primary triggers of the observed responses at P14. The increased Pdcl levels were attributed to the expression of a truncated Pdcl protein by the PhLPs transgene. Additionally, the upregulation of Gnb1 was consistent with PhLPs forming a stable tertiary complex with TRiC preloaded with nascent Gβ_1_[33].

By P18, the levels of 34 proteins, including five confirmed TRiC^et^ interactors, were significantly elevated (Fig 5D). The most upregulated proteins were the G-protein beta subunits Gnb2 and Gnb3, neither of which is typically expressed in rods at significant levels. These subunits closely resemble Gnb1, a component of rod-specific transducin, a heterotrimeric G protein involved in visual signaling. Given that PhLPs inhibits the TRiC-mediated folding of nascent Gnb1, it is plausible that the upregulation of Gnb2 and Gnb3 represents a compensatory response to transducin deficiency.

Additional upregulated TRiC^et^ interactors included histone-binding and chromatin remodeling proteins (Hdgfl2 and Hdac1) and the DNA-binding telomerase regulator Ylpm1. When analyzed as a whole, the upregulated proteins at P18 fell into four major functional groups. The largest group comprised transcriptional regulators and histones (e.g., Macroh2a2, Hdgfl2, H2ac4, H2ac6, Eif2b3, H2ac7, H2ac8, H2ac11, H2ac13, Hist1h2an, Hist1h2ao, Hist1h2ap, H2ac25, Hist1h2af, H2ac12, H2ac15, H2aj, H2ac1, Ddx3y, Ylpm1, Hnrnpul1, Eif2b3, and Hdac1). Three smaller groups included proteins involved in cytoskeletal regulation (e.g., Radil, Macf1, Dock7, and Camk2b), proteostasis maintenance (Psmd6 and Psmd13), and mitochondrial function (Pgam5 and Atp5f1e). Additional upregulated proteins included Prpsap1, which is involved in nucleotide biosynthesis, and Osbpl5, which participates in lipid transport. These findings suggest that by P18, the primary adaptive response of rod cells to the PhLPs transgene involves transcriptional regulation and chromatin remodeling.

Gene set enrichment analysis (GSEA) of the entire dataset revealed distinct effects of the PhLPs transgene at different time points after its expression. By P14, PhLPs predominantly suppressed recycling endosome, a subcompartment of the Golgi apparatus, and mitochondrial membrane while activating the proteasome and peptidase complexes (Fig 5E). At P18, the most strongly suppressed components included mitochondria, photoreceptor outer segment, and the cilium, whereas activation extended beyond the proteasome to include the translation initiation complex, nucleosome, DNA packaging complex and chromatin.

### Analysis of Proteins Within the PhLPs-TRiC Complex

We next examined how PhLPs influence the protein interactions of TRiC. To do this, we isolated the PhLPs-TRiC complex via FLAG-tagged PhLPs and analyzed its protein composition using LC-MS/MS (Sup. Table IV). The same protocol was applied as for TRiC^et^, with the analysis conducted at the proteomics facility of Duke University. Data from eight pulldowns performed on retinas from PhLPs and wild-type controls were averaged, and significantly enriched proteins that were also present in TRiC^et^ pulldowns were identified. To compare protein abundance, the mean intensity of the eight TRiC subunits in each dataset was calculated, and the intensity of each target protein was normalized to this value. The resulting normalized mean intensities were used to identify proteins that were significantly different between the PhLPs-TRiC and TRiC^et^ complexes (Fig 6).

**Fig. 6.**
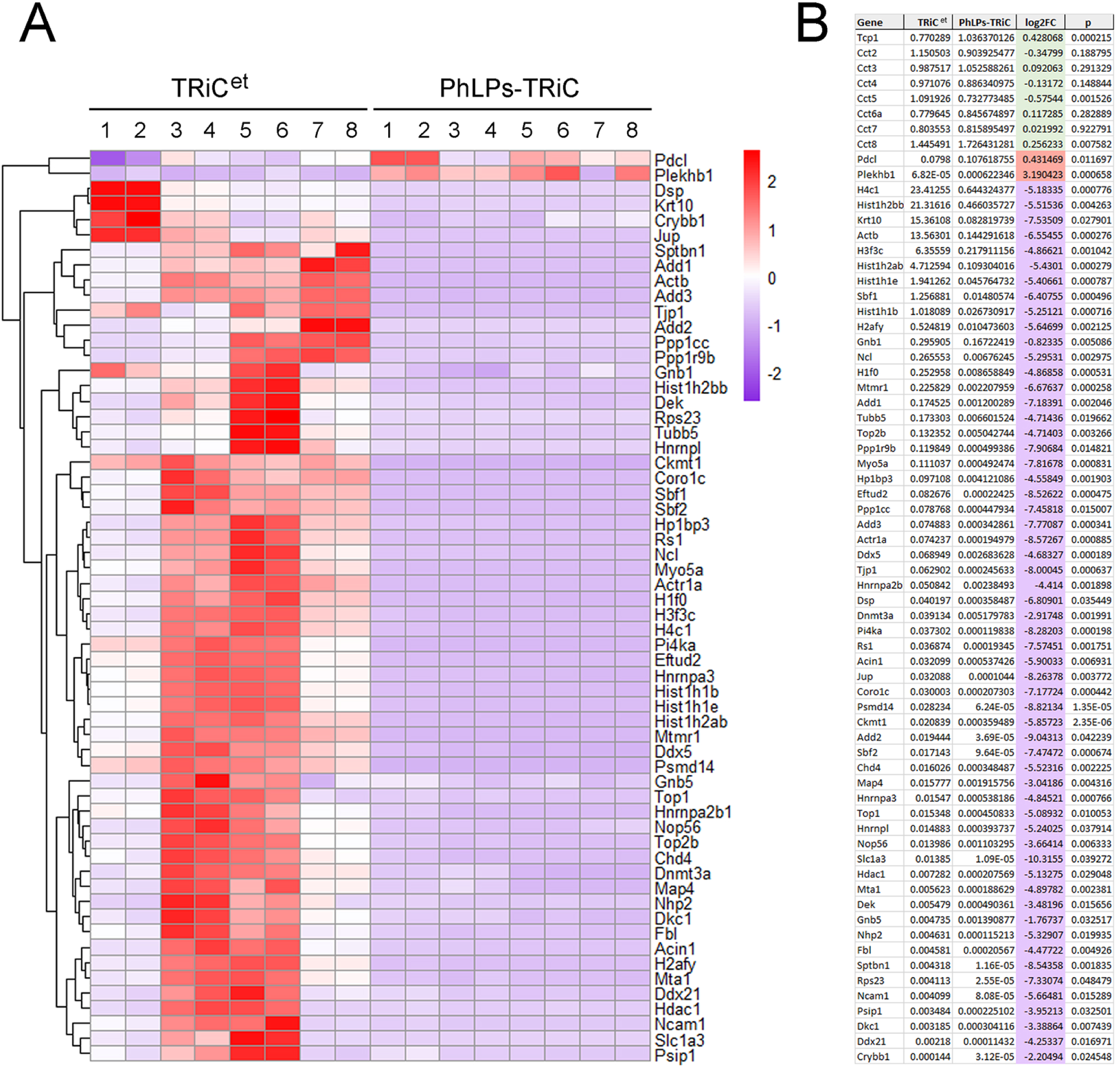
PhLPs transgene suppresses protein interactions of TRiC^et^. ***A***. Heatmap of relative protein abundance in the 900 kDa band in isolations from PhLPs and TRiC^et^ retinas, with z-scores indicating the relative normalized amount of protein within the PhLPs-TRiC complex compared to the TRiC^et^ complex (red: increased; blue: decreased). ***B***. Identified proteins (genes) ranked based on the numerical values of the log₂ fold change (log2FC) color-coded as red: increased, blue: decreased. The eight chaperonin subunits are in green.

In addition to the eight core subunits of the TRiC complex, 60 proteins were significantly enriched in both the PhLPs-TRiC and TRiC^et^ datasets. However, only two proteins—Pdcl and Plekhb1—were more abundant in the PhLPs-TRiC complex, while all other TRiC interactors were markedly reduced. The increase in Pdcl was expected, given that PhLPs is a splice isoform of Pdcl. The second enriched protein, pleckstrin homology domain-containing family B member 1 (gene: Plekhb1, protein: PKHB1), was increased by ninefold. Plekhb1 is known to be expressed in ciliated sensory neurons that utilize heterotrimeric G protein signaling, such as rod photoreceptors, though its specific function remains unclear[37–39]. Plekhb1 is an integral membrane protein that is not folded by TRiC. Interestingly, it has been shown to interact with the transducin Gβ_1_γ_1_ dimer[38], which is folded and assembled by TRiC and Pdcl[40,41]. This suggests that *Plekhb1* may play a role in regulating Gβγ biosynthesis.

All other identified proteins, including known and potential TRiC substrates such as Actb, Hdac1, Psmd14, Tubb5, Gnb1, and Gnb5, were significantly reduced in the PhLPs-TRiC complex. This finding provides further evidence that PhLPs suppresses TRiC-mediated protein folding in rod photoreceptors.

### Changes of Metabolic Profiles in the Retinas of PhLPs mice

Many TRiC interactors identified in our proteomic screen were enzymes, including two enzymes involved in glucose and nucleotide metabolism − Tpi1 and Uckl1− whose levels were significantly reduced in the retinas of PhLPs mice at postnatal day 14 (Fig 5B). Given the central role of TRiC in proteostasis, we hypothesized that its deficiency might extend beyond protein folding to broader metabolic disruptions in rod photoreceptors. To test this, we performed targeted metabolomics covering major metabolic pathways including glucose, nucleotide, amino acids and fatty acid metabolism[30]. In addition to PhLPs mice (treated group) and wild type 129E mice (control group), we included another group as disease control, composed of homozygous knock-in mice carrying the P23H mutation in the rhodopsin gene (*Rho*^P23H/P23H^). Both PhLPs and Rho P23H mice undergo progressive rod degeneration with similar onset and severity, but the underlying pathogenic mechanisms differ. Thus, the Rho P23H group served as a disease control to distinguish metabolic changes specifically attributable to TRiC dysfunction from those more generally associated with retinal degeneration.

Multivariate analysis of metabolomics data with PLSDA clearly separated treated from control retinas, demonstrating the overall difference in retinal metabolome between the control and treated groups (Fig 7A). Across 155 metabolites reliably detected in the retina, 37 and 51 showed statistically significant differences in the PhLPs and Rho P23H groups, respectively, compared to wild type controls (Fig 7BC, Sup. Tables V, VI). In PhLPs retinas, metabolic disruption was particularly evident in energy homeostasis and mitochondrial pathways (Fig 7D). We detected reductions in ATP, NAD, and NADH, accompanied by decreases in glycolytic intermediates (glucose, glucose 6-phosphate, 2-phosphoglyceric acid), TCA cycle metabolites (fumarate, α-ketoglutarate), and multiple acylcarnitines (octanoyl-, propionyl-, oleoyl-, isobutyryl-, and butyrylcarnitine). Together, these patterns suggest impaired glycolytic flux, reduced oxidative phosphorylation, and defective mitochondrial fatty acid oxidation. The loss of acylcarnitines, in particular, points to an underutilization of fatty acids as energy substrates, which may compromise the high bioenergetic demands of photoreceptors.

**Fig. 7.**
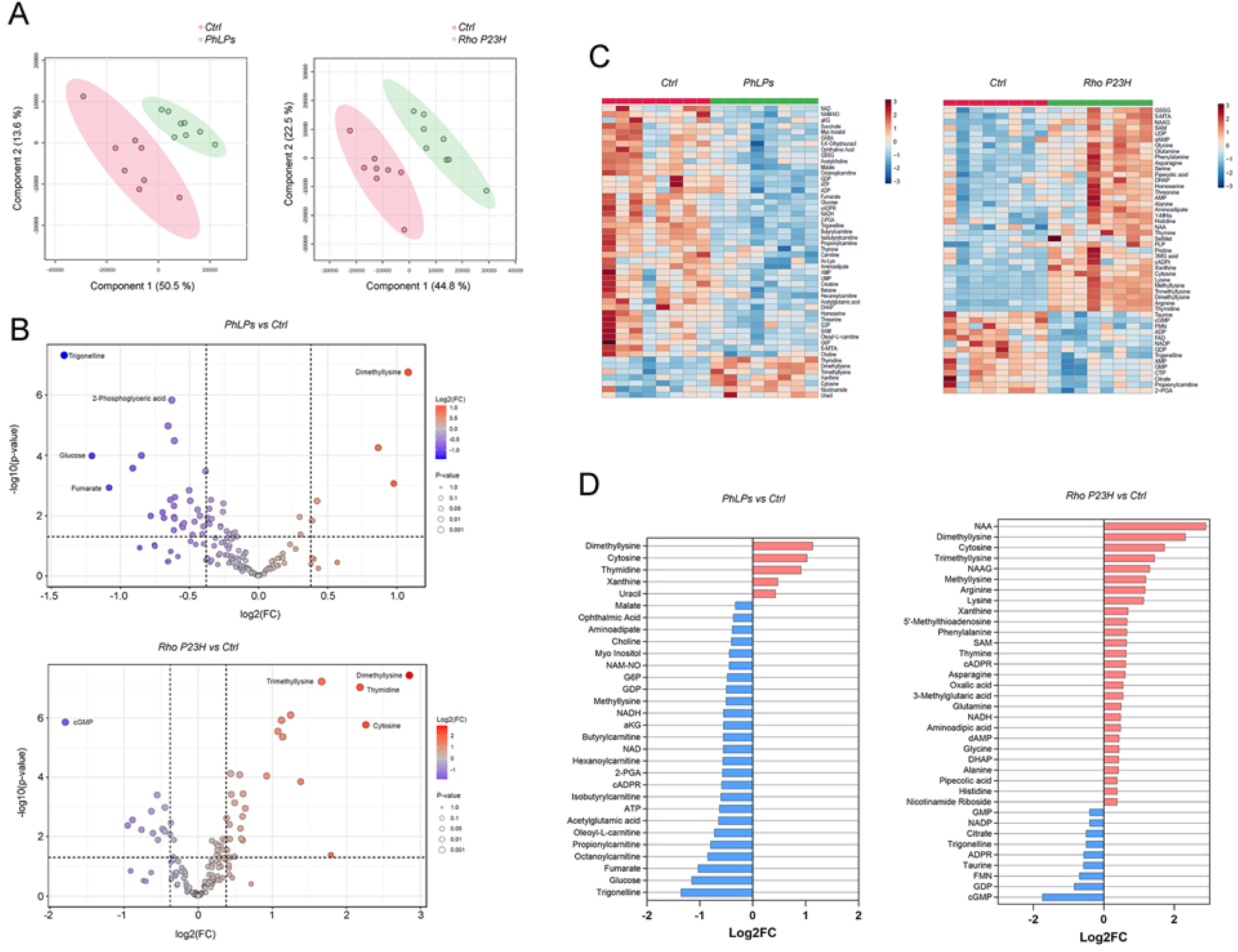
Metabolic changes in the retina of PhLPs and Rho P23H mice 14-days after birth ***A***. Partial Least Squares Discriminant Analysis (PLS-DA) scores plot illustrating group separation among WT, PhLPs, and Rho P23H retinas. Axes correspond to the first two latent variables (LV1 and LV2; % variance in Y explained shown on each axis). Ellipses indicate the 95% confidence regions for each group. ***B*.** Volcano plots of detected metabolites comparing either PhLPs or RhoP23H retinas to wild type 129E controls (n = 4 mice per group, 8 retinas analyzed independently). The x-axis shows log2 fold change (log2FC); the y-axis shows –log10(P value). The horizontal dashed line marks P = 0.05 (–log10P = 1.3). Significance was assessed by two-tailed t-tests (q < 0.05). ***C***. Heatmaps of the top 25 significantly altered metabolites for each comparison (left: PhLPs vs WT; right: RhoP23H vs WT). Rows represent metabolites, columns represent individual retinas. Values are row-scaled Z-scores, and both rows and columns were hierarchically clustered to emphasize tissue-specific patterns. **D.** Identified metabolites ranked based on the numerical values of the log₂ fold change (log2FC) color-coded as blue: decreased, red: increased (p<0.05, |log2FC|>0.4).

Beyond energy metabolism, TRiC deficiency also impacted NAD⁺ signaling pathways. Reduced cyclic ADP-ribose implies dysregulation of NAD⁺ metabolism, possibly reflecting overactivation of PARPs in response to cellular stress, further depleting NAD⁺ pools. This could exacerbate the energy crisis by limiting redox balance and sirtuin-mediated metabolic regulation. The decline in acetylglutamic acid, a critical urea cycle cofactor, suggests additional disturbances in nitrogen disposal and amino acid catabolism. Interestingly, these TRiC-dependent metabolic signatures were not mirrored in the Rho P23H model, despite comparable photoreceptor degeneration. Instead, Rho P23H retinas showed elevated levels of amino acids and methylation-related intermediates (Fig. 7D), pointing toward compensatory anabolic or stress-response pathways rather than energy collapse. In addition, deregulation of retinal cGMP level commonly observed in retinal degeneration models was readily detectable in Rho P23H group, but not in PhLPs group (Fig 7D).

Taken together, these results suggest that TRiC loss leads to a metabolic “energy crisis” in photoreceptors, characterized by collapse of glycolysis, TCA cycle activity, and fatty acid oxidation. This contrasts with the metabolic remodeling seen in Rho P23H retinas, underscoring the unique role of TRiC in safeguarding energy metabolism. We speculate that impaired proteostasis in PhLPs retinas may directly interfere with the folding and stability of key metabolic enzymes, thereby coupling chaperone failure to energetic collapse. Such TRiC-linked metabolic vulnerability could represent a previously unrecognized driver of photoreceptor degeneration.

### Integrated Proteomic–Metabolomic analysis identifies drivers of metabolic dysregulation in PhLPs Retinas

While our proteomic and metabolomic analyses independently revealed disruptions in protein homeostasis and energy metabolism, these layers of data only provide partial perspectives. To capture a more complete picture, we next performed an integrated analysis, combining proteomic and metabolomic datasets to identify protein drivers that may underlie the observed metabolite alterations using DIABLO. Such integrative approaches are particularly powerful because proteomic changes do not always translate linearly into metabolic outcomes, and metabolites can be influenced by both direct enzymatic regulation and broader cellular stress responses. By linking these data types, we aimed to identify the molecular nodes where TRiC dysfunction intersects with metabolic collapse in rod photoreceptors.

For this analysis, proteomic features were filtered based on fold change, while metabolomic features were prioritized by their opposite directionality compared to the Rho P23H disease-control group across five randomly selected replicates. This strategy enabled us to disentangle TRiC-specific metabolic alterations from generic degeneration-associated changes. The integration revealed a highly specific set of ten proteins (Rab10, Anxa1, Tuba1a, Txndc9, Uckl1, Wdtc1, Tubb4a, Hspa1a, Hspa1b) that co-varied with ten metabolites in the PhPLs group (Fig 8, Sup Table VII). Among these protein–metabolite pairs, two metabolites glucose and aminoadipic acid emerged as strongly driven by TRiC deficiency. Notably, glucose levels showed a direct correlation with Rab10 abundance, implicating defective Rab10-dependent GLUT4 trafficking in impaired glucose uptake in rods, a mechanism well-established in other cell types[42]. This provides a plausible link between TRiC dysfunction, disrupted proteostasis, and energy starvation in photoreceptors. In contrast, the relationship between Anxa1 and aminoadipic acid was less clear, lacking a known biochemical or signaling connection, but may suggest unexplored roles for annexins in amino acid homeostasis.

**Fig. 8.**
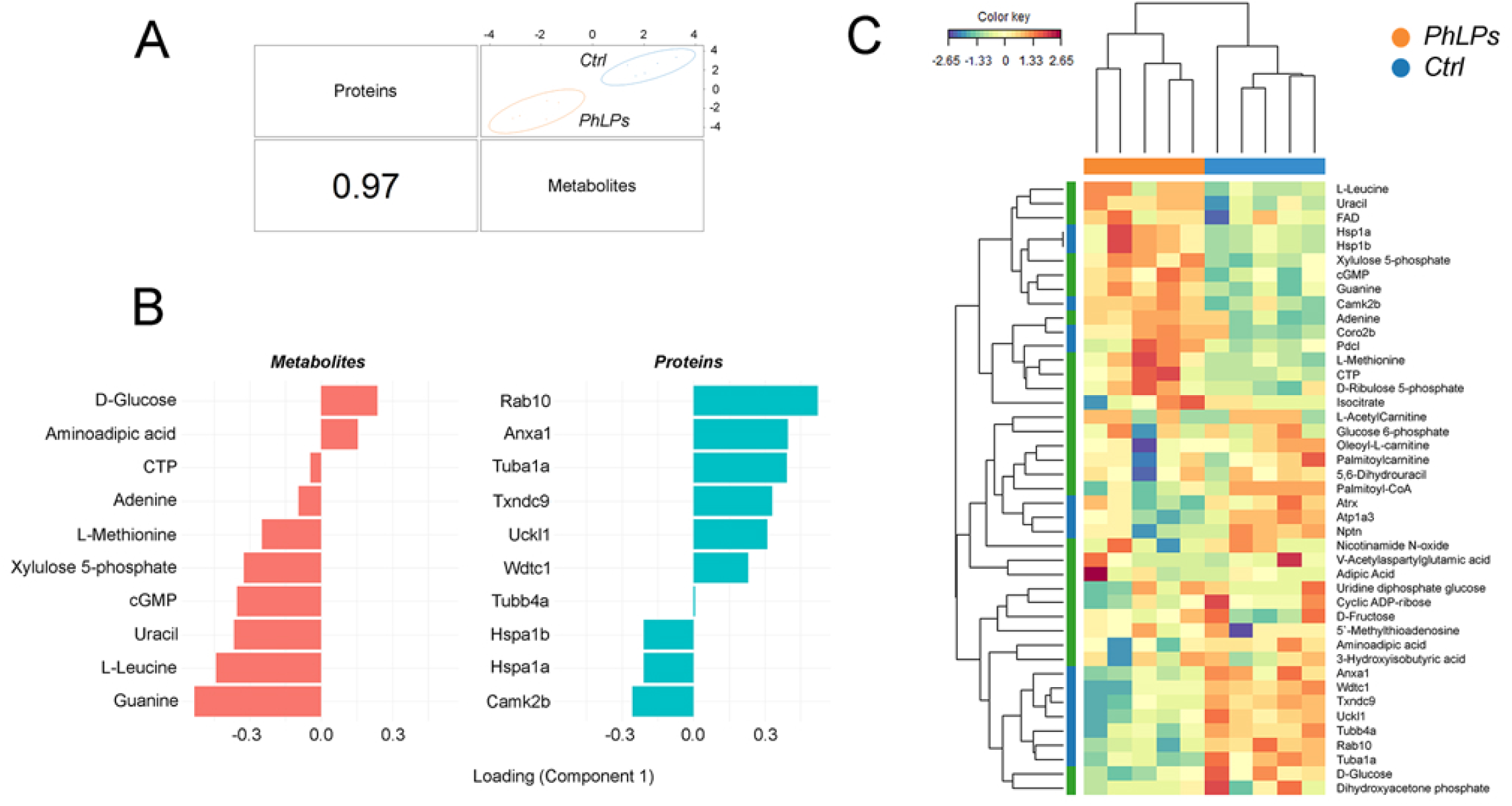
DIABLO-based integration uncovers TRiC-dependent molecular signatures in PhLPs retinas 14-days after birth. ***A***. Multiblock sparse PLS-DA (sPLS-DA) on proteomics and metabolomics datasets. Each sample is plotted on its first latent component for each block; points are colored by condition (wild type 129E control vs PhLPs). Ellipses denote 95% confidence regions. The correlation between the first components of the two blocks is shown (bottom left; r = 0.97) ***B***. Loadings plot highlighting the top proteins and metabolites (highest absolute loading weights) that discriminate wild type 129E control and PhLPs groups in the DIABLO model. ***C*.** Clustered Image Map (CIM) of the selected features across both omic blocks, proteomics and metabolomics. Cell colors represent pairwise correlations between features from DIABLO analysis (red = positive, blue = negative|). Variables with mean loading weight ≈ 0 were omitted.

This integrated omics analysis highlights how TRiC dysfunction triggers a cascade of metabolic stress centered on glucose utilization and mitochondrial bioenergetics. Importantly, this signature was not recapitulated in Rho P23H retinas, where metabolic changes likely reflect downstream degenerative processes rather than primary energetic defects. Thus, the integration of proteomic and metabolomic data helped us to uncover essential TRiC-specific molecular drivers such as Rab10 that would not have been evident from single-omic analysis alone. Collectively, these findings support a model in which TRiC safeguards photoreceptor metabolism by maintaining the stability of key trafficking and enzymatic proteins.

### Effect of Substrate Dominance on TRiC-Mediated Protein Folding

To evaluate the *in vivo* effect of a dominant TRiC substrate, we generated a perpetually unfolded mutant of Gβ_1_. The final step in the folding of nascent Gβ_1_ involves dimerization with the γ subunit [43]. To disrupt this process, we inserted alanine residues at positions 8 and 12 of myc-tagged Gβ_1_, disrupting the coiled-coil interaction with Gγ[44] (Fig 9A). This construct, termed myc-Gβ_1_^AA^, was co-expressed with HA-tagged Gγ_1_ in HEK293 cells, with wild-type myc-Gβ_1_ serving as a control. Both proteins were readily detectable in whole cell extracts; however, only myc-Gβ_1_ was present in HA pulldown assays, indicating that myc-Gβ_1_^AA^ failed to bind HA-Gγ_1_ (Fig 9B). We next compared the ability of myc-Gβ_1_ and myc-Gβ_1_^AA^ to interact with TRiC in HEK293 cells co-transfected with tcp-1α^et^. The 900 kDa TRiC complex was isolated via FLAG pulldown followed by blue native gel electrophoresis, excised, denatured with SDS-containing buffer, and analyzed by Western blotting using anti-myc antibodies (Fig 9C). We found that myc-Gβ_1_^AA^ was enriched ∼1.8-fold compared to wild-type myc-Gβ_1_ in TRiC^et^ complexes (Fig 9D), suggesting that the mutant persists in an unfolded state and preferentially binds TRiC (Fig 9E).

**Fig. 9.**
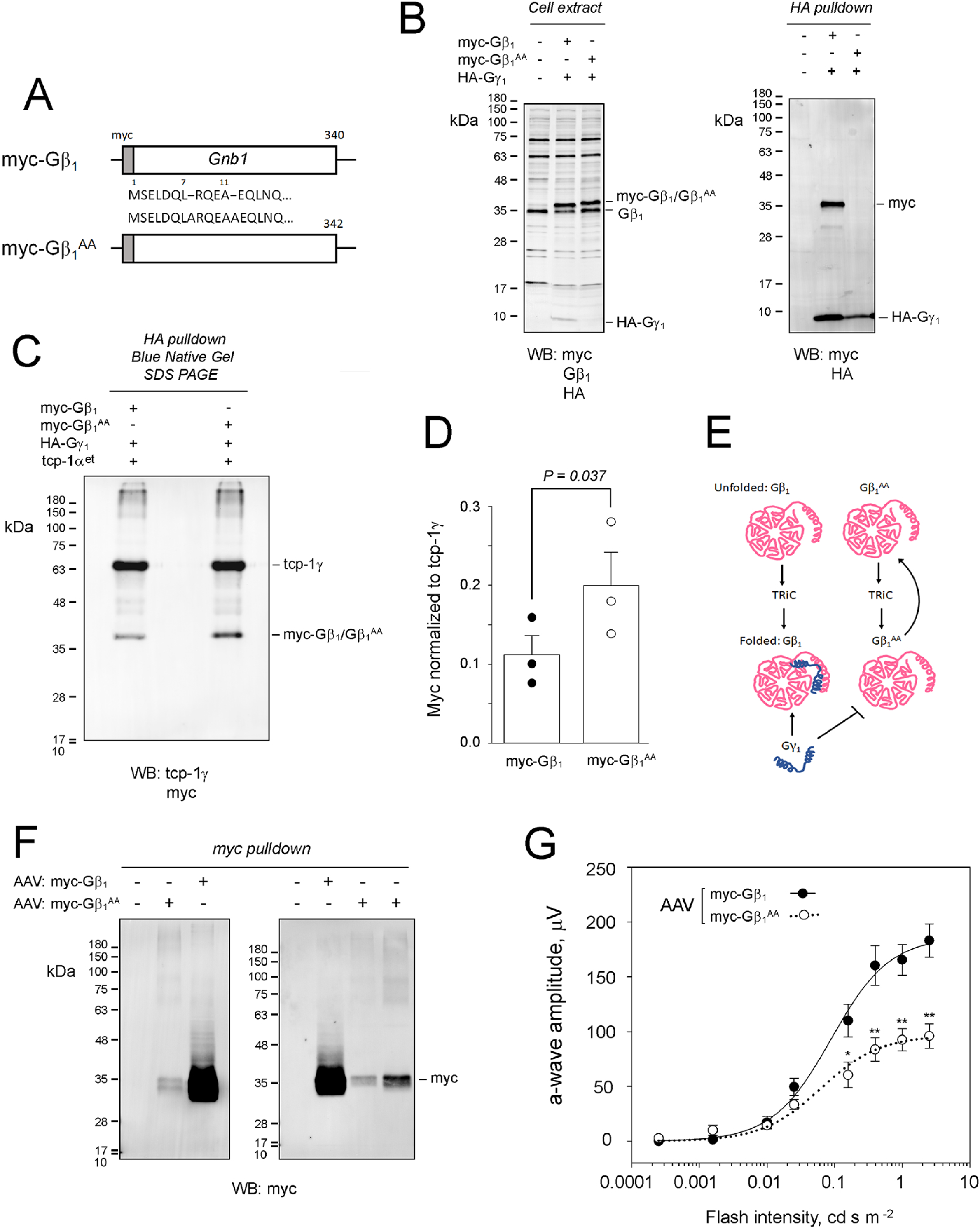
Perpetually unfolded Gβ_1_ mutant increasingly binds TRiC and is cytotoxic for rods. ***A***. Epitope-tagged rod transducin β subunit (gene: *Gnb1*; protein: Gβ_1_), myc-Gβ_1_, and its mutant with Ala^8^ and Ala^12^ insertions, myc-Gβ_1_^AA^. ***B***. Western blot analysis of HEK293 cells co-transfected with epitope-tagged rod transducin γ subunit (gene: *Gng1*; protein: Gγ_1_) designated as HA-Gγ_1_ and myc-Gβ_1_ or myc-Gβ_1_^AA^: whole cell extract (left), anti-HA pulldown (right). ***C***. Western blot analysis of the 900 kDa band from Blue Native Gel. Anti-FLAG pulldown was conducted in HEK293 cells co-transfected with tcp-1α^et^, HA-Gγ_1_ and myc-Gβ_1_ or myc-Gβ_1_^AA^. ***D***. Normalized amounts of myc-Gβ_1_ and myc-Gβ_1_^AA^ in the 900 kDa band; bars are mean value with SE, P-value determined by t-test, n=3. ***E***. Cartoon of Gγ_1_ completing the TRiC-mediated folding of Gβ_1_, disrupted in Gβ_1_^AA^ mutant. ***F***. Western blot analysis of anti-myc pulldown from the retina. Wild type 129E mice received subretinal injection of AAV- myc-Gβ_1_ and AAV - myc-Gβ_1_^AA^ and their retinas collected and analyzed 1-month post-injection. ***G***. Comparison of visual responses of mice that received subretinal injection of AAV - myc-Gβ_1_^AA^ (white circles) and AAV- myc-Gβ_1_ (black circles) 4-months post-injection. Electroretinographic responses were recorded using a UTAS BigShot (LKC Technologies) rodent ERG system. The amplitude of elicited ERG a-wave is plotted as a function of flash intensity (SEM, n= 5 (myc-Gβ_1_) n=6 (myc-Gβ_1_^AA^)). Each dataset was fitted with a simple rectangular hyperbola with two parameters. The significance was determined by t-test and Mann-White rank sum test, with P value <0.05 (*) and <0.01 (**).

To assess its effect *in vivo*, we delivered myc-Gβ_1_^AA^ to mouse rod photoreceptors via subretinal injection of AAV vectors, using myc-Gβ_1_ as control. One-month post-injection, myc signal levels were compared. Because myc-Gβ_1_^AA^ was barely detectable in whole retina extracts, we used myc pulldown to confirm expression, which was substantially lower than that of wild-type myc-Gβ_1_ (Fig 9F). Longitudinal electroretinography showed a significant reduction in a-wave amplitude in mice treated with AAV: myc-Gβ_1_^AA^ at 4 months post-injection, indicating rod dysfunction (Fig 9G). Thus, the myc-Gβ_1_^AA^ mutant, unable to bind Gγ_1_, was less stable and more cytotoxic in rod photoreceptors than wild-type Gβ_1_. Next, we sought to determine how unfolded Gβ_1_ influences TRiC protein interactions. Because AAV vectors result in patchy transduction in mouse rods, they were not suitable for proteomic analysis. Instead, we crossed TRiC^et^ mice with Gγ_1_⁻/⁻ mice[19]. Given that Gγ_1_ is the predominant γ subunit in mouse rods, its absence is functionally equivalent to the AA mutation in producing unfolded Gβ_1_. As reported for AAV: myc-Gβ1AA-treated rods, Gγ_1_^⁻/⁻^ photoreceptors undergo progressive degeneration[19,45]. We used this genetic model to analyze TRiC interactions by LC-MS/MS using our standard approach. Data from six TRiC^et^ pulldowns in Gγ ^⁻/⁻^ and wild-type retinas were averaged, and significantly enriched proteins identified. The same analysis was performed for TRiC^et^ vs. wild-type samples. For normalization, we calculated the mean intensity of the eight TRiC subunits in each dataset and used this value to normalize the intensity of each target protein. These normalized average intensities were used to identify significant differences in TRiC^et^; Gγ_1_^⁻/⁻^ vs. TRiC^et^ datasets (Fig 10A, Sup Table VIII). Using this approach, we identified 10 proteins with increased association, including four previously validated TRiC^et^ interactors (Cenpv, Pdcd5, Gnb1, and Pdcl) and six new interactors (Gfap, Ica, Tprl, Hnrnpm, Syne1, and Actr1b (Fig 10C). Notably, Gβ_1_ (Gnb1) showed a 2.2-fold increase, consistent with the enrichment observed for myc-Gβ_1_^AA^ (Fig 9D), indicating its unfolded state. The increased TRiC-Gβ_1_ interaction likely stimulated binding of Pdcl, a co-chaperone mediating the folding of Gβ_1_ [32]. Among 52 proteins with decreased association were 12 known TRiC interactors and substrates (Atg16l1, Wdr31, Txndc9, Tubb5, Wdtc1, Tuba4a, Tubb2a, Hnrnpll, Iars, Gnb3, Gnb5, and Plekhb1). This group also included three bona fide TRiC substrates from the tubulin family: Tubg1, Tubb3, and Tuba1a (Fig 10D). Together, these data demonstrate that unfolded Gβ_1_ increasingly occupies TRiC, displacing other substrates when it fails to properly fold due to the absence of Gγ_1_. This competitive binding may underlie Gβ_1_ cytotoxicity and could, in theory, amplify the detrimental effects of misfolding mutations in other TRiC substrates. For example, when tubulin folding in rods is impaired, microtubules in cilium and elsewhere may become unstable or dysfunctional. Accordingly, the increased interaction of TRiC with the cytoskeleton- and stress-related proteins, such as Cenpv[46], Gfap[47], Pdcd5[48], Syne1[49], and Actr1b may reflect an attempt to stabilize or reconfigure microtubules. Likewise, impaired folding of Atg16l1 may compromise the formation of autophagosome[50], which may be further exacerbated by the increased binding of Tiprl, a positive regulator of mTORC1 inhibiting autophagy[51] essential for rod photoreceptor survival[52]. The suppression of autophagy may prevent degradation of TRiC via lysosome-autophagy pathway observed in the HeLa cells treated with cytochalasin D, resulting in TRiC inactivation by unfolded actin[53].

**Fig. 10.**
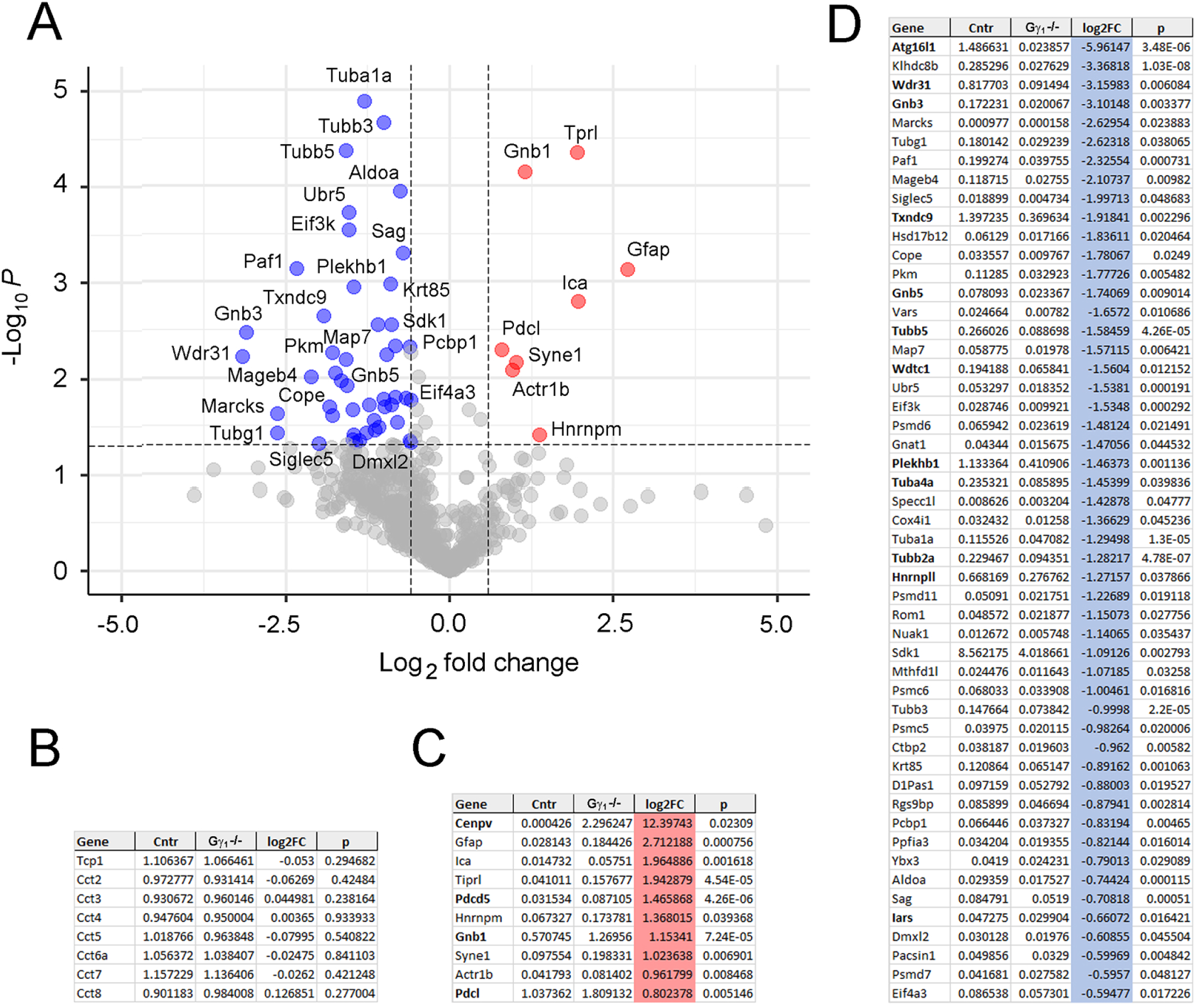
Perpetually unfolded Gβ_1_ affects protein interactions of TRiC^et^ in rods. ***A***. Volcano plot of differential protein levels in the 900 kDa band isolated from the retina. TRiC^et^ mice of Gγ_1_^-/-^background and TRiC^et^ mice of wild type 129E backgrounds are compared at one month of age. The x-axis represents the log₂ fold change (log₂FC) and the y-axis shows (–log₁₀(p-value), with p < 0.05 and |logFC| > 1.5 inclusion criteria. The data were collected at the Proteomic Facility at the Duke University. ***B***. Normalized levels of the eight subunits of TRiC, the numerical values of the log₂ fold change (log2FC), and p-value in the compared groups. ***C***. Normalized levels of proteins increased in TRiC^et^ preparations from Gγ_1_^-/-^ retinas (red) ranked based on log₂FC. ***D***. Normalized levels of proteins decreased in TRiC^et^ preparations from Gγ_1_^-/-^ retinas (blue) ranked based on log₂FC. Independently identified TRiC^et^ interactors (Sup. Table II) are in bold font.

## Conclusions

Our comprehensive proteomic analysis of TRiC interactors in rod photoreceptors, leveraging both data-dependent acquisition (DDA) and data-independent acquisition (DIA) approaches, revealed 226 candidate proteins. Despite deploying complementary DIA and DDA workflows on identically prepared material, the overlap between datasets and with published interactomes remained limited. We interpret this not as experimental noise but as an inherent limitation of current discovery proteomics: each method detects a different subset of *bona fide* targets while missing others. Several canonical TRiC substrates reproducibly emerged across our screens, strengthening confidence in the core client set, but some notable negatives, including actin, remain unexplained. The absence of actin, an established TRiC substrate, may reflect one or more factors: limited turnover in mature rods, peptide detectability issues, or competitive occupation of TRiC by dominant clients such as unfolded Gβ_1_ or tubulins. These possibilities highlight an important unresolved biological and technical question. Bioinformatic analyses of the combined interactome revealed that a surprisingly large number of TRiC interactors had RNA-recognition motif and RNA-related functions, implicating TRiC in RNA processing and splicing pathways in rod photoreceptors.

Characterization of the effect of the short PhLPs splice isoform on TRiC, building on our previous studies, confirmed that PhLPs toxicity is mediated through TRiC. Our data demonstrate that PhLPs-bound TRiC loses its ability to engage its other substrates, while the mutation reducing PhLPs binding to TRiC make it less cytotoxic. However, the physiological *in vivo* role of the PhLPs isoform remains elusive. Functional studies using the PhLPs-induced TRiC loss-of-function model demonstrated that tubulins are the dominant TRiC clients whose folding is acutely sensitive to TRiC inhibition, explaining the severe disruption of rod outer segment biogenesis. However, the most striking and reproducible finding of this study was the TRiC-linked collapse of energy production in PhLPs retinas, spanning glycolysis, TCA intermediates, acylcarnitines, and NAD(H), a phenotype absent in a degeneration-matched rhodopsin mutant control. This argues for a TRiC-specific metabolic vulnerability rather than a generic consequence of photoreceptor loss and points to a mechanistic axis connecting proteostasis failure (Tpi1/Uckl1 destabilization) with impaired glucose handling via Rab10/GLUT4 trafficking.

Previously, we reported that suppression of TRiC disrupts the assembly of the BBSome complex, a finding supported by reduced levels of Bbs2, Bbs5, and Bbs7 in the PhLPs retinas, and proposed that these proteins are folded by TRiC[18]. These proteins are core components of the BBSome complex, whose dysfunction causes Bardet-Biedl syndrome (BBS), a ciliopathy characterized by retinal dystrophy[54]. Although Bbs2, Bbs5, and Bbs7 were not identified as TRiC interactors in our current interactome analysis, Bbs1 was identified as a candidate TRiC substrate significantly enriched in the 900 kDa fraction. It is plausible that the 65 kDa Bbs1 protein is a TRiC substrate and that its deficiency could underlie incomplete BBSome assembly in PhLPs-expressing rods[18]. This idea is supported by clinical evidence, as mutations in Bbs1 are the most common cause of BBS[55,56]. Furthermore, knock-in mice carrying the M390R mutation in Bbs1 develop defects in both rod outer and inner segments, leading to retinitis pigmentosa-like retinal degeneration[57]. However, the more rapid degeneration observed in the PhLPs mouse model suggests the destabilization of the cytoskeleton resulting from TRiC suppression [58–60] profoundly exacerbates the pathological phenotype caused by BBSome deficiency.

Finally, our study provides evidence that genetic mutations stabilizing TRiC clients in their non-native, unfolded states can have cytotoxic consequences. We propose the following hypothetical mechanism to explain this effect. Under normal physiological conditions, the concentration of nascent, non-native proteins is very low, and our data are consistent with this: in our TRiC^et^ pulldowns, the relative abundance of client proteins was extremely low. For example, in the Duke dataset, 125 out of 131 significant TRiC interactors were present at a molar ratio of 0.025 or less relative to TRiC^et^ (mean: 0.0142 ± 0.006; median: 0.001). This suggests that TRiC is typically present in vast excess over its substrates, preventing competition among clients for TRiC binding. However, this delicate balance would be disrupted if the concentration of a single misfolded substrate were greatly increased. Such an overload could sequester TRiC and hinder the folding of other essential clients. Through this mechanism, misfolding mutations in TRiC substrates may induce global proteostasis collapse, amplifying cellular dysfunction.

In summary, our integrated proteomic analyses reveal that TRiC not only assembles canonical substrate complexes but also uniquely engages retinal-specific client proteins involved in phototransduction and cellular stress responses, underscoring essential roles of TRiC for maintaining proteostasis in rod photoreceptors. We identified novel TRiC clients, proposed mechanisms linking TRiC dysfunction to retinal degeneration, and highlighted the potential for substrate-specific misfolding mutations to trigger broader cellular collapse. These insights enrich our understanding of TRiC’s biology and function in the retina but also suggest that TRiC may play a critical role in other neurodegenerative diseases driven by proteostasis imbalance. Future work aimed at modulating TRiC activity or enhancing substrate folding capacity could offer promising therapeutic strategies for these conditions.

## Acknowledgments

We thank Dr. Paolo Fagone and the Biochemistry Viral Core for generating AAV vectors.

This work is supported by U.S. National Institutes of Health grants R01EY030056 to W.D., R01AG0641888 to D.S., R01 EY005722 to N.S., R01019665, R01 EY030050 to M.S., R01 EY031324, R01 EY032462 and the Retina Research Foundation grant to J.D..

IDeA National Resource for Quantitative Proteomics is supported by U.S. National Institutes of Health grant R24GM137786.

The authors declare no competing interests.

During the preparation of this manuscript, the author, Maxim Sokolov used Chat GPT to improve readability and English. After using this tool, Maxim Sokolov reviewed and edited the content as needed and takes full responsibility for the content of the published article.

